# Spindle: Scalable Indexing of Spatial-Omics Data for Fast Querying

**DOI:** 10.64898/2026.11.12.766059

**Authors:** Souvik Ghosh, Stavros Sintos, Anrin Chakraborti, Hirak Sarkar

## Abstract

Spatial transcriptomics (ST) has unlocked new avenues for biological discovery by jointly capturing gene expression and spatial context. ST datasets are typically stored as gene expression matrices paired with spatial coordinates, supporting queries based on expression levels at individual locations. However, characterizing how genes vary together requires representations of relationships between genes, such as covariance matrices computed over spatial neighborhoods. These gene–gene co-expression patterns capture coordinated biological variation and can reveal spatially localized gene programs. As datasets grow to atlas scale, efficiently searching for such patterns becomes a central computational bottleneck due to the combination of high dimensionality, extreme sparsity, and variability in gene expression across large numbers of spatial regions. Such searches require repeated access to second-order statistics, motivating a compact, searchable representation of spatial co-expression.

We address this problem with Spindle, a platform-agnostic indexing algorithm for efficiently searching spatial gene co-expression patterns. Spindle exploits the approximate block-diagonal structure of gene–gene covariance matrices over spatial regions, in which covariances between blocks are small in magnitude, to selectively store informative components while compactly representing recurring covariance patterns. This yields a compressed yet searchable index suitable for atlas-scale spatial datasets. Across multiple real-world biological datasets (87,000–817,000 cells), Spindle indexes are 113–197 times smaller than the whole covariance matrices. A complete query, including query preprocessing, takes 2.8–28 ms on one CPU core, 5.1–16 times faster than a scan over the whole covariance matrices, and the tiles it returns recover a median of 81% of the exact near neighbors. In breast cancer, the regions retrieved by covariance search closely match the cell-type composition of the query (median composition difference 0.15, compared with 0.54 for random regions). Spindle supports both partial covariance queries and queries specified as gene lists, as well as retrieval across measurement platforms.

**Availability and implementation:** Spindle is implemented in Python with compiled C search kernels and is available as open-source software under GPL v3 at https://github.com/sarkar-lab/spindle.

## 1 Introduction

Spatial transcriptomics (ST) measures gene expression at spatial locations within intact tissue sections. Public ST collections now encompass hundreds of datasets and millions of cells across diverse tissues and biological conditions [1, 2], generated by platforms with different measurement resolutions and gene coverage: imaging-based assays such as Xenium measure targeted panels of hundreds to thousands of genes at cellular or subcellular resolution, whereas sequencing-based assays such as Visium measure transcriptome-wide expression in spots approximately 55 μm in diameter [3, 4]. As these collections grow, researchers need efficient ways to retrieve regions that resemble a region of interest, much as sequence databases support searching with BLAST [5] or cell atlases support searching with a query cell [6–8].

Existing ST resources offer limited support for this form of region-level retrieval. SpatialDB [9] and STOmicsDB [10] provide datasets for download and visualization, while SOAR [11] supports queries based primarily on individual-gene expression, often within pre-annotated cell types. None provides a general mechanism for retrieving tissue regions by the similarity of their multigene expression patterns, so researchers must download and preprocess datasets before performing their own comparisons.

Many tissue programs are characterized not only by the expression levels of individual genes, but also by the coordinated expression of groups of genes [12, 13]. Recurring programs such as epithelial–mesenchymal transition, angiogenesis, and immune evasion are observed across tumor types [14], and spatially resolved co-expression can reveal structures such as stem-cell niches [15]. A gene–gene covariance matrix summarizes this co-expression within a spatial region and can distinguish regions with similar average expression but different patterns of coordinated expression; in image analysis, region covariance matrices are likewise used as descriptors for detection and matching [16]. Covariance-based descriptions of spatial neighborhoods have been used to identify cellular niches [17], but, to our knowledge, no existing method retrieves tissue regions by the similarity of their gene–gene covariance matrices.

Efficiently searching these matrices presents two challenges. First, a covariance matrix over *p* genes has *p*(*p* + 1)*/*2 distinct entries (49,141 for a 313-gene panel), so storing and comparing them across thousands of regions is expensive. Second, similarity comparisons should respect the matrix structure; for positive-definite matrices, the log-Euclidean distance provides a computationally convenient metric on the symmetric positive-definite (SPD) cone [18–20]. Generic nearest-neighbor indexes [21, 22] could be applied to vectorized matrix logarithms, but each region would contribute *p*(*p* + 1)*/*2 entries, and each query would still require the logarithm of a whole *p* × *p* matrix.

Here, we present Spindle, a compact index for similarity search over gene–gene covariance matrices of spatial tiles (Fig. 1). Rather than storing an independent representation of every covariance matrix, Spindle exploits recurring patterns of gene–gene covariance across tiles. It groups tiles into covariance niches and, within each niche, identifies a common gene ordering under which the covariance matrices are approximately block diagonal, so that the logarithms of the blocks, rather than of whole matrices, can be shared across tiles. For each niche, a layered directed acyclic graph (DAG) stores shared representative block logarithms, each tile is a path through it, and a distance-budgeted depth-first search prunes paths whose lower bounds exceed the budget, while returning every tile within it. Spindle supports complete queries (a tile’s covariance matrix over all genes), partial queries (a covariance matrix over a subset of genes, searched without reconstructing the missing entries), and gene-list queries (a template covariance matrix constructed from a list of genes). As an exhaustive reference, Spindle-naive stores the block logarithms of every tile and ranks all tiles using the same block distance.

**Fig. 1:**
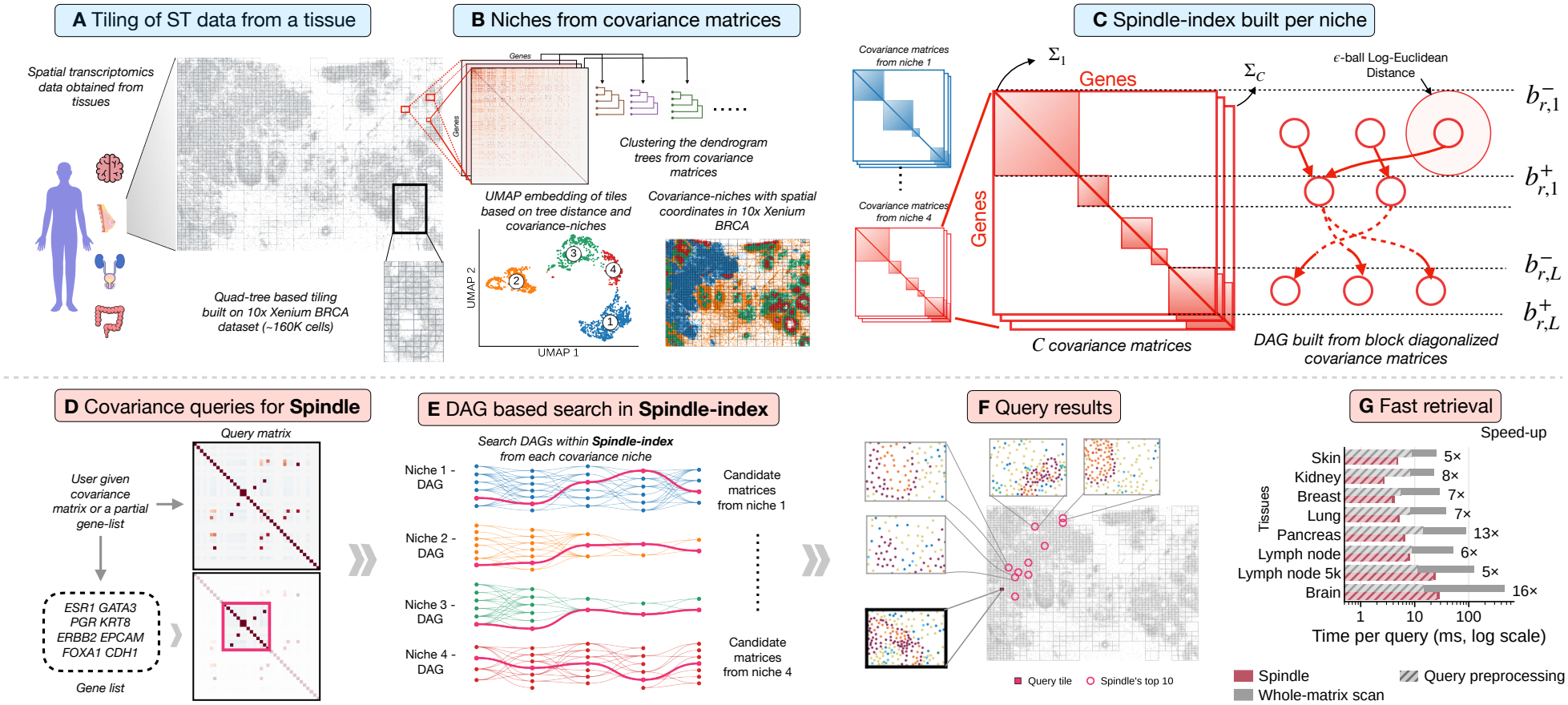
Overview of Spindle. **A** A Xenium breast-cancer dataset of approximately 160,000 cells, partitioned by a quadtree into tiles of at most 200 cells. **B** Covariance niches: tile covariance matrices are converted to gene-correlation dendrograms and grouped by dendrogram similarity. Left, UMAP [23] of tiles based on tree distance, colored by niche; right, the niches in the tissue. **C** Index construction for one niche. Left, example covariance matrices from niches 1 and 4 in their gene orders and block partitions. Middle, the *C* covariance matrices Σ_1_, …, Σ_*C*_ of one niche *r*; dotted lines mark the first and last genes, 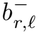 and 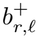, of block *ℓ*. Right, the layered DAG, with one layer per block, nodes as circles (log-Euclidean balls of radius *ϵ*), and edges between nodes in consecutive layers that share a tile. **D** Query: a user-provided covariance matrix (top) or a gene list giving a template covariance matrix (bottom; magenta outline). **E** DAG search in each niche; paths are tiles colored by niche, and paths within the distance budget (magenta) give the candidate tiles. **F** Query tile (magenta square) and the first 10 tiles returned by Spindle (magenta circles). **G** Time per complete query on one core, on a log scale, for Spindle and a whole-matrix scan: query preprocessing (hatched) followed by the search (solid); labels give the speed-up of Spindle. Panels **C–E** are schematic; **G** shows the mean over five random splits.

We evaluate Spindle on eight Xenium datasets to assess index size, query time, and retrieval accuracy for complete and partial queries. Using paired Xenium and Visium sections from the same breast tumor, we test whether covariance-based retrieval transfers across measurement platforms and preserves cell-type composition. We further evaluate gene-list queries derived from 24 curated tissue signatures to assess whether retrieved regions exhibit the corresponding expression patterns. These experiments examine whether a space-frugal covariance index can support efficient retrieval while preserving biologically relevant similarities.

## 2 Results

### 2.1 Overview of Spindle

Given an ST dataset, Spindle first tiles the tissue using a quadtree [24, 25], with each tile containing at most a specified number of cells (e.g. 200 for 10x Xenium). This produces smaller tiles covering densely populated regions and larger tiles covering sparse regions (Fig. 1A). Spindle then computes a gene–gene covariance matrix for each tile and groups them into “covariance niches” (Section 3.2) by a similarity metric [26, 27] based on the order of genes (Fig. 1B). Within each niche, Spindle partitions the ordered gene list into contiguous blocks so that the covariance matrices are approximately block diagonal: for each tile, covariance entries within the *diagonal blocks* are expected to have larger absolute values than those outside the blocks (Supplementary Fig. S1). Spindle compares tiles using the block distance *d*_*B*_ (Eq. (3)) defined as the log-Euclidean distance [18] between the corresponding diagonal blocks (Fig. 1C). The exhaustive base-line Spindle-naive uses the same niche-specific gene orders and diagonal blocks to compute *d*_*B*_ between the query and every indexed tile, returning tiles ranked by increasing distance.

For each niche, Spindle builds a layered DAG where each layer corresponds to a diagonal block. To avoid repeatedly storing similar principal submatrices, Spindle clusters the submatrices corresponding to the same diagonal block across tiles within each layer (Fig. 1C). Each node of the DAG stores a representative principal submatrix and a radius *ϵ* defining a ball that contains the corresponding submatrices of all tiles assigned to that node (Fig. 1C). Each tile is represented by a path through the DAG, with one node per layer. Although the niches and blocks are constructed for efficient indexing, they also reflect the biology of the underlying tissue architecture: in breast cancer, one niche outlines the DCIS ducts and their myoepithelial layer, another the invasive tumor, and a third the stroma and its immune infiltrate, and several blocks are enriched for known T-cell, fibroblast and proliferation programs (Supplementary Fig. S3). In practice, for efficient computation, Spindle stores the logarithm of the representative principal submatrices rather than the raw matrix (Eq. (S5)).

Spindle provides a general-purpose framework for retrieving spatial regions whose gene–gene covariance patterns are similar to those of a given query. It supports three types of queries (Fig. 1D): (1) a **complete query**, the covariance matrix of a tissue region over all genes in the index, taken from the indexed dataset or from another dataset; (2) a **partial query**, a covariance matrix over only a subset of these genes, that is, the submatrix of a complete query on those genes; and (3) a **gene-list query**, a list of genes from which Spindle constructs a template covariance matrix (Eq. (7)) and searches it as a partial query.

For each niche-specific DAG, Spindle first reorders the query covariance matrix according to the niche’s gene order and extracts the principal submatrices corresponding to its diagonal blocks. It then performs a greedy search on the DAG to find the node whose representative principal submatrix is closest to the query’s principal submatrix at each layer. The best match establishes a distance budget, after which a depth-first search explores only paths whose lower bounds on *d*_*B*_ fall within that budget (Fig. 1E). Candidates from all niches are ranked by their query distance, so retrieved tiles can come from anywhere in the dataset (Fig. 1F), and the number of retrieved tiles depends on the query. For a gene set *S*, Spindle uses the partial block distance 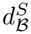 on the same niches and blocks, organizing the genes within each block into dyadic intervals (section 3.5). Across eight Xenium datasets, Spindle answers complete queries, including query preprocessing, 5.1–16 times faster than a whole-matrix scan on one core (Fig. 1G).

### 2.2 Setup and evaluation

We evaluate Spindle on eight public Xenium datasets of skin melanoma [28], non-diseased kidney [29], breast cancer [3], lung cancer [30], pancreatic cancer [31], lymph node [32], lymph node profiled with the 5k panel [33], and brain cancer [34] (Fig. 2A; Supplementary Table S1). For each dataset, we hold out a random 10% of the tiles as queries and index the rest, repeating with five random seeds; unless otherwise specified, we report means across the five splits.

**Fig. 2:**
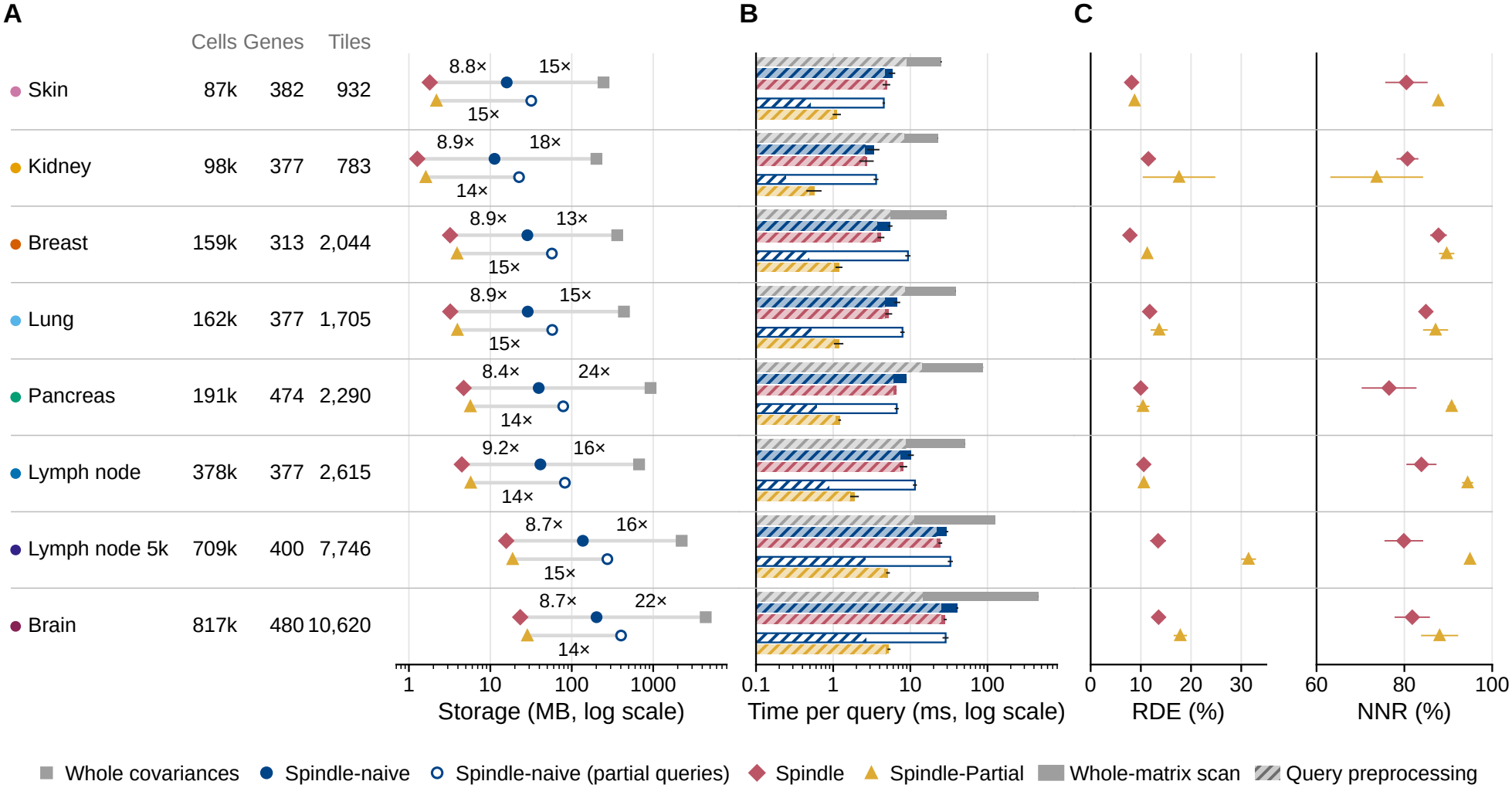
Spindle achieves compact indexing, fast search, and accurate retrieval across eight Xenium datasets. **A** Index storage on a log scale; the table lists the cells, genes, and tiles of each dataset. Upper row, whole covariance matrices, Spindle-naive, and Spindle for complete queries; lower row, Spindle-naive and Spindle-Partial for partial queries. Labels give the fold reduction from Spindle-naive to Spindle (upper row, left; lower row) and from the whole matrices to Spindle-naive (upper row, right). **B** Time per query on one core, on a log scale: query preprocessing (hatched), followed by the search (solid), for a whole-matrix scan, Spindle-naive, and Spindle (complete queries) and Spindle-naive and Spindle-Partial (partial queries). Bars and error bars show the mean and standard deviation across five random splits of the median time over the first 200 held-out queries (all, if fewer); the hatched part shows the median time of query preprocessing. **C** RDE (left) and NNR (right) at *c* = 5% of the index size for complete (diamonds) and partial (triangles) queries; mean and standard deviation across five random splits. The NNR axis begins at 60%.

We evaluate retrieval accuracy using relative distance error (RDE) and near-neighbor recall (NNR), assess biological similarity using cell-type composition difference, and report query times for computational efficiency.

#### Retrieval accuracy

We measure accuracy against the exact ranking of Spindle-naive, using 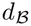 for complete queries and 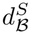 for partial queries; within a niche, *d*_*B*_ orders tiles nearly as the whole-matrix log-Euclidean distance does (Spearman correlation 0.942–0.955 in the three largest breast-cancer niches; Supplementary Fig. S2). Let 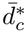 and 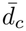 denote the mean distances to the *c* nearest index tiles and to the first *c* tiles returned by Spindle, respectively, and let 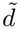 denote the median distance to all index tiles. The relative distance error (RDE) is

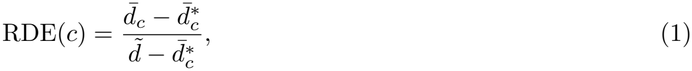

which equals 0 when the returned tiles have the same mean distance as the exact nearest neighbors and 1 when their mean distance equals the median distance to all index tiles. We set *c* to 5% of the index size, or to the number of returned tiles if Spindle returns fewer. We call an index tile a near neighbor of the query if its distance is at most 1.05 times that of the nearest index tile; near-neighbor recall (NNR) is the fraction of these near neighbors that appear among the first *c* returned tiles.

#### Cell-type composition difference

To assess the biological similarity of retrieved tiles to the query tile, we measure their *cell-type composition difference*. We compare these differences against a background distribution of composition differences between the query tile and randomly selected index tiles to assess whether retrieved tiles have more similar cell-type compositions than expected by chance.

#### Partial query evaluation

For a partial query, we restrict the covariance matrix of a held-out tile to |*S*| ∈ {4, 8, 16, 32} genes that are consecutive within one block, taking the gene order and blocks from the niche of the tile’s nearest index tile. As baselines, we complete the missing entries and search Spindle-naive, either by *padding* the missing genes with zero covariance or by *imputing* them from the mean covariance matrix of the niche whose mean is closest to the query.

Query times are measured on a single core unless stated otherwise (Supplementary Table S2).

### 2.3 Spindle enables fast, accurate co-expression search with space-frugal indexes

Spindle indexes are 113–197 times smaller than the full covariance matrices and about nine times smaller than Spindle-naive indexes (Fig. 2A). Much of this reduction comes from the block representation itself, which already makes Spindle-naive 12.7–23.6 times smaller than the whole matrices, and sharing block representatives across tiles reduces storage further. The gain is largest for larger gene panels, since whole-matrix storage grows quadratically with the number of genes whereas each block contains at most 100 genes, and the index grows approximately linearly with the number of tiles (Supplementary Fig. S4 and Table S1).

These compact representations also make queries fast (Fig. 2B). On a single core, a complete query takes 2.8–28 ms with Spindle compared with 23–456 ms for a whole-matrix scan, a 5.1–16-fold speedup. These times include query preprocessing, which computes the required matrix logarithms and accounts for 90–95% of the total query time for Spindle (Supplementary Table S3). The search itself takes 0.17–2.9 ms, 48–152 times faster than the scan. Much of the speedup comes from comparing diagonal blocks rather than whole matrices, while the DAG further reduces the number of tile comparisons. For partial queries, Spindle-Partial takes 0.58–5.3 ms, 4.0–7.8 times faster than Spindle-naive, and its search alone is 38–56 times faster (Supplementary Table S4).

Spindle supports parallelization of both query preprocessing and search (Supplementary Fig. S5). During preprocessing, matrix logarithms can be computed independently across niches and diagonal blocks; during search, each niche can be processed independently. A whole-matrix scan can also compare tiles in parallel, but its query preprocessing requires computing the logarithm of a single dense *p* × *p* matrix, which took 4.9– 14 ms and showed little improvement with additional cores. As cores are added, both approaches therefore approach a floor, and that of Spindle is much lower: on 32 cores, a complete query takes 0.46–2.0 ms with Spindle and 7.3–43 ms with a whole-matrix scan, 13–25 times faster with Spindle.

The efficiency gains come with modest retrieval accuracy (Fig. 2C). For complete queries, the median RDE across datasets is 11.0%, and the first *c* returned tiles (*c* = 5% of the index tiles) recover a median of 81% of the query’s near neighbors; partial queries achieve similar accuracy on most datasets. The ranking error arises from using distances to node representatives rather than to individual tiles’ exact block logarithms (Supplementary Fig. S6B). In contrast, completing the missing entries of a partial query and searching Spindle-naive returns tiles much farther from the query than those of Spindle-Partial (Supplementary Fig. S6D,E).

### 2.4 Spindle retrieves tissue regions with similar cell-type composition across platforms

Spindle retrieves tissue regions with similar cell-type composition across ST platforms, despite differences in spatial resolution and gene coverage. We evaluate this on serial sections of the same breast tumor profiled with Xenium and Visium [3], registered to common coordinates with SIID [35] and restricted to the 307 genes shared by both panels (Fig. 3A). We partition the Xenium section into tiles and overlay the same boxes on the Visium section, so that each Visium tile has a co-located Xenium tile, and use each tile of one platform as a query against an index of all tiles of the other.

**Fig. 3:**
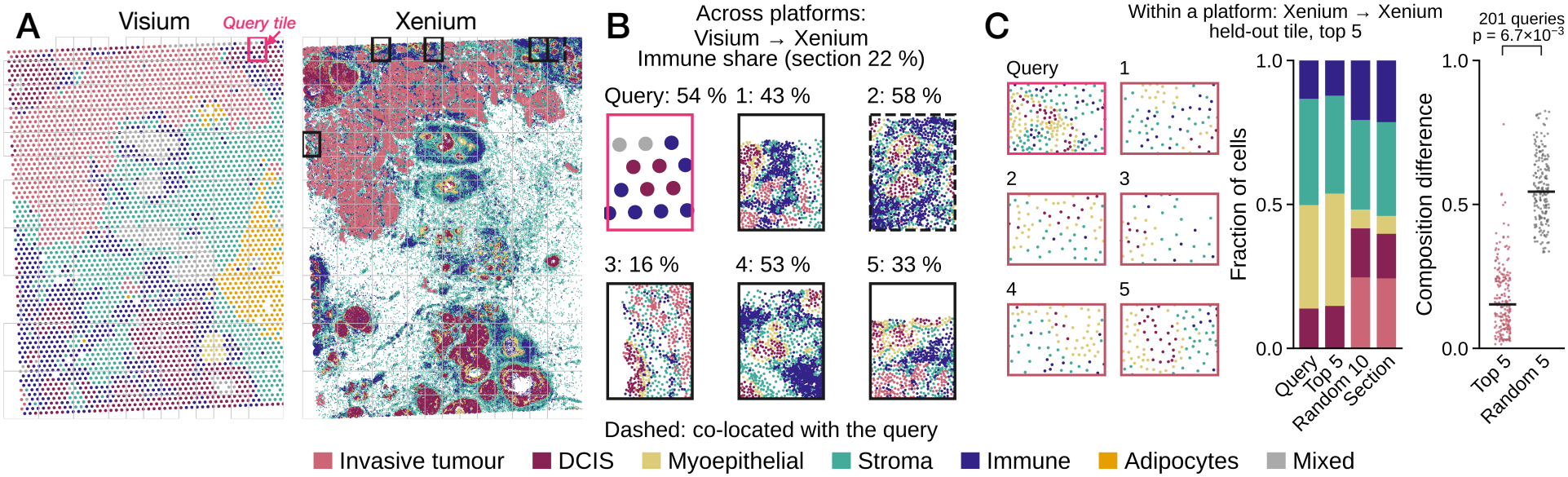
Spindle retrieves compositionally similar tissue regions within and across platforms. **A** Serial breast-cancer sections profiled with Visium (left; 55 *μ*m spots) and Xenium (right; cells), colored by cell-type annotation. Gray boxes are the tiles overlaid on both sections (at most 2,000 Xenium cells each); the magenta box marks the Visium query of **B**, and black boxes the five Xenium tiles retrieved by Spindle (dashed, co-located). **B** The Visium query and its first five retrieved Xenium tiles after the global adjustment of query logarithms. Labels give the immune-cell proportion of each tile and of the full Xenium section; the dashed tile is co-located with the query. **C** Within-platform retrieval with held-out Xenium tiles. Left, a query tile and its first five retrieved tiles. Middle, cell-type composition of the query, the five retrieved tiles (pooled), 10 random index tiles (averaged over 200 draws), and the full dataset. Right, composition difference between each held-out query and its first five retrieved tiles or five random index tiles (averaged over 200 draws); horizontal lines mark medians, and the *p*-value is from a one-sided Wilcoxon signed-rank test.

The covariance logarithms of the two platforms differ systematically (Supplementary Fig. S7A). Before searching, we therefore apply an unsupervised global adjustment that shifts and rescales the logarithm of each query’s covariance matrix to match those of the index tiles (section A.7). After this adjustment, retrieved tiles have a cell-type composition similar to that of their queries. The tumor fraction of a query and the mean tumor fraction of its first 10 retrieved tiles are correlated for queries from either platform (Pearson’s *r* = 0.81 for Visium queries and 0.73 for Xenium queries; Supplementary Fig. S7B,C). For example, a Visium query rich in immune spots retrieves Xenium tiles enriched for immune cells, including its co-located tile (Fig. 3B).

Spindle also retrieves compositionally similar regions within a platform (Fig. 3C). Across held-out breast-cancer queries, the median composition difference to the first five retrieved tiles is 0.15, compared with 0.54 for five random index tiles, and the retrieved tiles are closer for almost every query (one-sided Wilcoxon signed-rank test [36]). Similarity in gene–gene covariance patterns can thus identify spatial regions with similar cell-type composition, both within a dataset and across measurement platforms.

### 2.5 Partial and gene-list queries retrieve biologically relevant regions across datasets

We evaluate whether partial and gene-list queries retrieve biologically relevant regions in three settings (Fig. 4): (i) partial queries from another dataset; (ii) partial queries from held-out tiles of the same dataset; and (iii) gene-list queries.

**Fig. 4:**
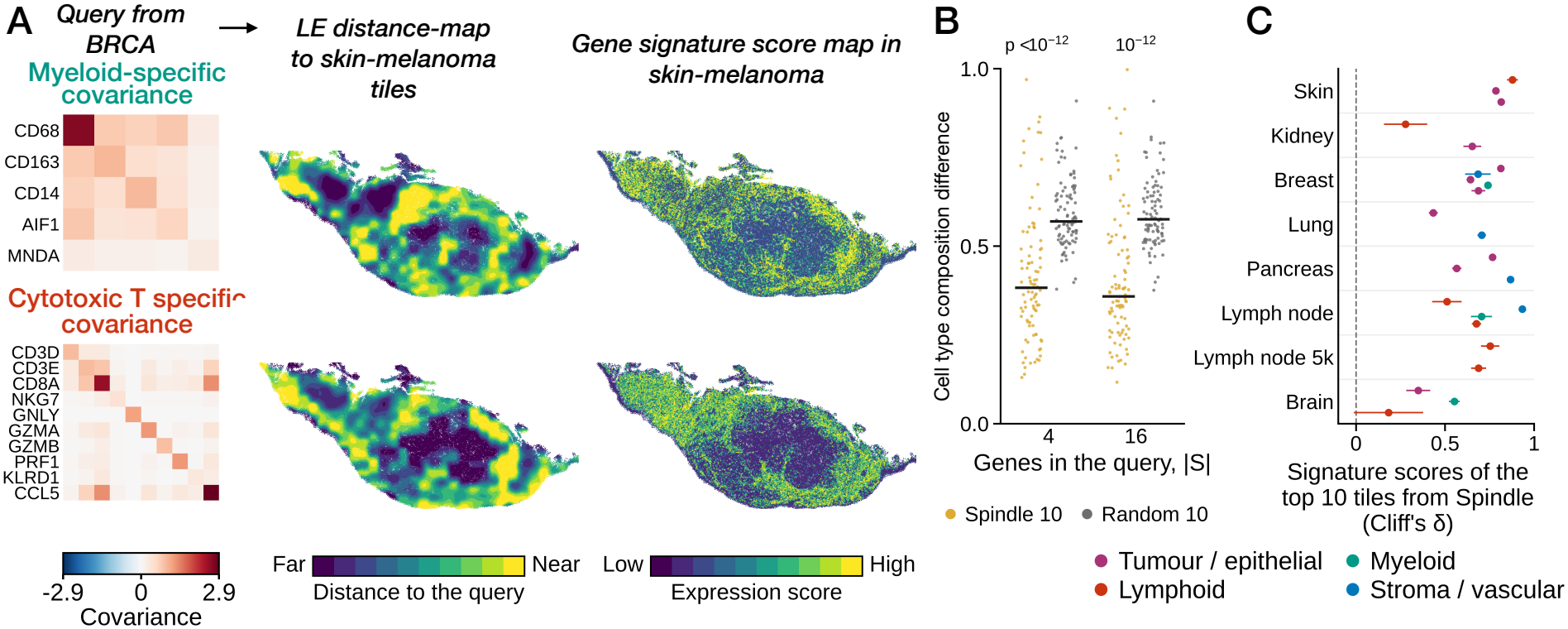
Partial queries and gene-list queries retrieve biologically relevant tissue regions. **A** Cross-dataset partial queries from breast-cancer (BRCA) tiles searched against the skin-melanoma index, for myeloid-cell (top) and cytotoxic T-cell (bottom) markers. Left, query covariance matrices. Middle, query closeness from the exact partial block distance 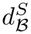 (log-Euclidean, LE), projected onto the tissue at tile centers and smoothed with a Gaussian kernel of one tile side. Right, mean z-scored expression of the same genes in each cell. Color scales show percentiles across cells in the main tissue. **B** Composition difference between each of the 93 held-out skin tiles, as a partial query over 4 or 16 genes, and the first 10 tiles returned by Spindle (yellow) or 10 random index tiles (gray); bars mark medians, and *p*-values are upper bounds from one-sided Wilcoxon signed-rank tests. **C** Cliff’s *δ* between the expression scores of up to 10 tiles retrieved by gene-list queries and those of the remaining index tiles, for 24 curated tissue signatures colored by class (tumor/epithelial, lymphoid, myeloid, stroma/vascular); mean and standard deviation across five random splits.

To test partial queries across datasets, we use two marker sets, five myeloid-cell genes and 10 cytotoxic T-cell genes. Each query takes its gene–gene correlations from the breast-cancer tile in which the marker genes are most strongly co-expressed (section A.8). We then compute the distance from each query to every tile of the skin-melanoma index with Spindle-naive and map it onto the tissue (Fig. 4A). Tiles close to the query lie in regions where the marker genes are highly expressed, and query closeness is positively correlated with the expression score of the marker genes (Spearman’s *ρ* = 0.45 for myeloid and 0.37 for cytotoxic T-cell genes).

For partial queries from held-out tiles of the same dataset, the first 10 retrieved tiles are closer in composition to the query tile than 10 random index tiles for most queries, with median composition differences of 0.38 against 0.57 for four-gene queries (Fig. 4B). The other seven datasets show the same trend with smaller differences (Supplementary Table S6).

Finally, for gene-list queries from 24 curated tissue signatures, we compare the expression scores of the retrieved tiles with those of the remaining index tiles using Cliff’s *δ* [37], where *δ* = 1 means that every retrieved tile scores higher than every remaining tile (Fig. 4C; Supplementary Table S7). Cliff’s *δ* is positive for all 24 signatures and at least 0.5 for 20 of them, indicating that a gene list alone, without a measured covariance matrix, retrieves regions in which the listed genes are highly expressed.

## 3 Methods

### 3.1 Background

We represent an ST dataset by *N* cells, with spatial coordinates *z*_*i*_ ∈ ℝ2 and raw gene-count vectors *x*_*i*_ over a gene panel *G* consisting of *p* genes. We partition the cells with a quadtree [24, 25], starting from the bounding box of all cells and recursively splitting each box at the midpoints of its sides until every box contains at most 200 cells. The resulting bounding boxes are referred to as tiles. Tiles with very low cell density, at the tissue edges and in debris, are discarded as outliers. For a tile *T*, let *X*_*T*_ ∈ ℝ|*T* |*×p* be its cell-by-gene count matrix. We compute a preprocessed covariance matrix Σ_*T*_ for tile *T* (see section B.1.3). By construction, Σ_*T*_ is a symmetric positive-definite (SPD) matrix. We build the Spindle index from a set of *n* tiles and write Σ_*m*_ for the covariance matrix of index tile *m* (section B.1).

Conventional Euclidean distance does not capture the geometry of symmetric positive-definite matrices because they form a non-Euclidean Riemannian manifold. The log-Euclidean distance addresses this limitation by mapping matrices to the space of symmetric matrix logarithms [18], providing a computationally efficient distance respecting the positive-definite structure. For an SPD matrix *A* = *V* diag(*λ*_*k*_)*V* ^⊺^, let log *A* = *V* diag log max(*λ*_*k*_, 10^−6^) *V* ^⊺^. The log-Euclidean distance between two SPD matrices is defined as

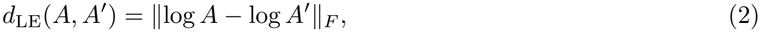

where ∥ · ∥_*F*_ is the Frobenius norm [18, 19].

Gene–gene covariance matrices are often approximately block diagonal after the genes are reordered, with small entries outside the blocks. We define a *block structure ℬ* = (*σ*; *B*_1_, …, *B*_*L*_), where *σ* is a permutation function over *G*, and *B*_*j*_ denotes a set of contiguous gene indices over *σ* that represents a block in a covariance matrix with *L* such blocks. A matrix is block diagonal under *ℬ* if all its entries outside the blocks *B*_1_, …, *B*_*L*_ are zero. For any two block-diagonal SPD matrices with the same block structure, their squared log-Euclidean distance decomposes additively into the sum of the squared log-Euclidean distances between corresponding blocks: 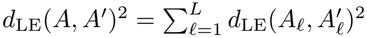 (see Supplementary Theorem S1). Spindle exploits this property by not using the covariances between blocks, so that the distance between two tiles is computed block by block.

Apart from making the pairwise distance calculation faster, the block structure *ℬ* itself is also biologically meaningful. Each block is a group of genes that are co-expressed within a tile. However, different groups of genes are co-expressed in different parts of a tissue, so the block structure varies across tiles. Therefore, using one block structure *ℬ* might not be ideal for every tile from biologically heterogeneous tissue. Spindle resolves this problem by grouping tiles with similar block structure and referring to them as covariance niches [17].

### 3.2 Covariance niche construction

Spindle partitions the tiles into covariance niches *N*_1_, …, *N*_*J*_, such that each niche contains tiles with similar gene covariance and shares one block structure 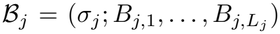. We write *n*_*j*_ = |*N*_*j*_|, *E*_*j*_ for the block-log dimension under *ℬ*_*j*_, and *j*(*m*) for the niche containing tile *m*.

To construct the niches, Spindle first embeds each tile’s covariance matrix Σ_*T*_ in up to 30 dimensions. It groups genes using average-linkage hierarchical clustering [26], with distance 1 − |*r*_*gg′*_ |, where *r*_*gg′*_ is the correlation between genes *g* and *g*′ in tile *T* . The resulting dendrogram defines cophenetic distances, given by the height at which each pair of genes first joins the same cluster [27]. These distances, collected in a fixed gene-pair order, are summarized by PCA across tiles [38] to obtain the tile embeddings (see section B.3.1). Finally, Leiden clustering [39] of the 10-nearest-neighbor graph of these embeddings, with the resolution increased until no niche contains more than 1,000 tiles, gives the niches.

The same dendrograms also give the block structure of each niche. Spindle applies average-linkage clustering to the mean cophenetic matrix of the tiles in a niche to construct a consensus dendrogram (see section B.3.3), whose leaf order defines *σ*_*j*_. We cut this dendrogram at the knee of its run-count curve (see Supplementary Fig. S8) into fine runs of at most 50 genes and merge adjacent runs into blocks of 15 to 100 genes by a dynamic programming algorithm that maximizes Σ _*ℓ*_ *W*_*j*_(*B*_*j,ℓ*_)*/*|*B*_*j,ℓ*_|2, where *W*_*j*_(*B*) is the sum of the squared mean correlations in the niche between distinct genes of *B* (section B.3 and Supplementary Fig. S8). This construction serves as a heuristic for finding a consensus block structure for each niche (see section B.3.3).

Within each niche, Spindle compares tiles using only the principal submatrices *A*[*B*_*ℓ*_] of the shared block structure *ℬ* = (*σ*; *B*_1_, …, *B*_*L*_), without using the covariances between blocks. Instead of the log-Euclidean distance of whole matrices, it therefore uses the *block distance* and the *partial block distance*. Formally, given a nonempty gene set *S* ⊆ *G*, let *S*_*ℓ*_ = *S* ∩ *B*_*ℓ*_. The block distance and the partial block distance between *A* and *A*′ are

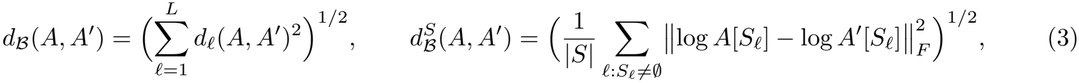

where 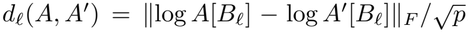. Thus, *d*_*ℬ*_ is the log-Euclidean distance between the block-diagonal parts of *A* and *A*′, normalized by 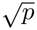, and ignores covariances between genes in different blocks; leaving them out lowers the distances between tiles of a niche by a nearly constant amount and hardly changes their order (Supplementary section A.1). For the partial block distance, each covariance matrix is restricted to *S*_*ℓ*_ before its logarithm is taken; consequently, a partial query requires covariances only for the queried genes. In practice, Spindle stores the upper-triangular entries of each block logarithm as a vector vec_*ℓ*_(*A*), scaled so that the Euclidean distance between two such vectors equals *d*_*ℓ*_ (section B.2).

### 3.3 Building the Spindle index

Within each niche, Spindle clusters the block logarithms of all tiles and organizes the resulting representatives in a layered directed acyclic graph (DAG). Formally, given a niche *N*_*j*_ whose tiles share the block structure 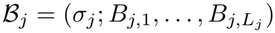, Spindle constructs the DAG *G*_*j*_ = (*V, F* ), where 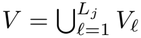 and *V*_*ℓ*_ is the set of nodes in layer *ℓ*. Each layer corresponds to one block, and each node *v* ∈ *V*_*ℓ*_ stores a representative block-log vector *μ*_*v*_ and a radius *r*_*v*_ (see Fig. 1C and section B.5).

In each layer, Spindle selects *K* representatives with a greedy approximation to the *K*-center problem (see Supplementary Problem S2). For each layer *ℓ*, farthest-first traversal [40] selects up to 256 of the layer’s blocklog vectors as candidate centers; for every *K*, the first *K* centers have a covering radius *ρ*_*ℓ*_(*K*) at most twice the optimal *K*-center radius. We allocate retained centers across layers by prioritizing the largest reduction in squared covering radius per unit of storage, weighted by block spread, with a storage budget of *β* = 0.10 times that of Spindle-naive for the niche (see section B.5 and Supplementary Eq. (S25)). Each tile *m* is assigned to its nearest retained center in every layer, denoted *a*_*ℓ*_(*m*). After construction, each node *v* = (*ℓ, k*), corresponding to the *k*-th retained center in layer *ℓ*, stores its center *μ*_*v*_, member set *M*_*v*_ = {*m* : *a*_*ℓ*_(*m*) = *k*}, and covering radius *r*_*v*_ (the *ϵ* in Fig. 1C).

### 3.4 Searching Spindle

Spindle searches with two kinds of query covariance matrices: a *complete query Q* over all genes and a *partial query Q*_*S*_ over a nonempty gene set *S* ⊆ *G*. For an index tile *m*, we define the query-to-tile distance as 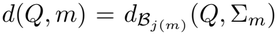 for a complete query covariance matrix *Q*, and 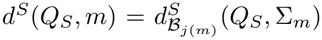 for a partial query *Q*_*S*_ over a nonempty gene set *S*. Here *j*(*m*) is the covariance niche containing tile *m*, and *ℬ*_*j*(*m*)_ is the block structure of that niche. For a query Spindle searches all the niches to find the best match. In practice, finding all matches might not be useful; as a result, Spindle solves a budgeted search problem defined below.

#### Problem 1

(Budgeted covariance search). Given a query and a distance budget *η* ≥ 0, let *δ*(*m*) = *d*(*Q, m*) for a complete query, or *δ*(*m*) = *d*^*S*^(*Q*_*S*_, *m*) for a partial query. The objective is to return every index tile *m* satisfying *δ*(*m*) ≤ *η*, ranked by increasing *δ*(*m*).

While searching the DAG, Spindle uses node-level lower bounds to prune paths that exceed a query-specific distance budget. For a query covariance matrix *Q* and niche *j*, let 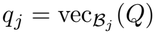, with *q*_*j,ℓ*_ denoting the entries for the block in layer *ℓ*. For each node *v* in that layer, we compute

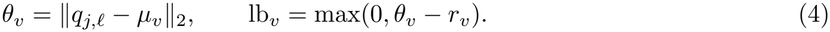

The node bounds combine along each path to give a lower bound on the distance to every tile following that path (Supplementary Lemma S1).

#### Path intersection

The search maintains the tiles that follow each path prefix *P* = (*v*_1_, …, *v*_*s*_) through the intersection 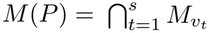 (Fig. 1E). Only extensions with nonempty intersections are explored, ensuring that every visited path contains tiles and each tile is returned at most once.

#### Probe, budget and search

A greedy probe in each niche selects, at each layer, the node with the smallest *θ*_*v*_ among those that share a tile with the path so far, giving a probe score 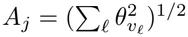. The best probe score sets a shared distance budget,

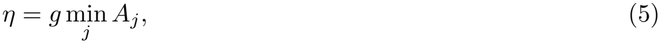

where *g* controls search breadth and defaults to 0.25. A depth-first search explores nodes in increasing order of lb_*v*_, pruning paths whose combined lower bound exceeds *η*. Tiles on completed paths are ranked across niches by the approximate score 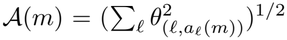. Tiles satisfying *d*(*Q, m*) ≤ *η* are ranked approximately using node centers as representatives (Supplementary Theorem S2). Proofs and algorithmic details are given in section B.6.

The exhaustive reference index, Spindle-naive, stores 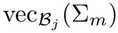 for every tile *m* in every niche *j*, requiring Σ_*j*_ *n*_*j*_*E*_*j*_ values, computes *δ*(*m*) for all index tiles and sorts them, thereby solving Problem 1 exactly. For partial queries, it additionally stores the block covariance matrices and computes log Σ_*m*_[*S* ∩ *B*_*j,b*_] for each nonempty block intersection at query time (sections B.4 and B.7.1). In contrast, Spindle stores only a fraction of these values, selects the budget *η* from the query, and returns all tiles within that budget ranked by an approximate score.

### 3.5 Partial and gene-list queries

To search partial queries, Spindle-Partial builds nodes as in Section 3.3 for the dyadic intervals of each block: sets of positions *I* = {*a*, …, *a* + 2^*k*^ − 1} within the block, numbered from zero in the niche’s gene order, with *a* ≡ 0 (mod 2^*k*^); a block of *p*_*b*_ genes contains fewer than 2*p*_*b*_ such intervals. For every interval, nodes and node assignments are built from the upper-triangular entries of log Σ_*m*_[*I*], with off-diagonal entries multiplied by 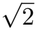 and the covariance matrix restricted to *I* before its logarithm is taken, using the same storage fraction *β*. The node-allocation gain divides the reduction in squared covering radius by the interval’s spread, rather than multiplying by it, so that intervals have more comparable relative approximation error. No edges are stored: the node assignments determine the valid extensions through the path intersection *M* (*P* ) (section B.7.2).

For a query over *S*, we process each niche independently. Within each block *B*_*j,b*_, we decompose each maximal run of consecutive positions of *S* ∩ *B*_*j,b*_, from left to right, into the fewest dyadic intervals (Supplementary Fig. S9). The resulting intervals *I*_1_, …, *I*_*s*_ form the layers of the search of Section 3.4, with node distances divided by 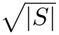 and *g* = 1 by default. The argument of Supplementary Theorem S2 guarantees that the search returns every tile within the budget under the piece distance over the intervals *I*_1_, …, *I*_*s*_

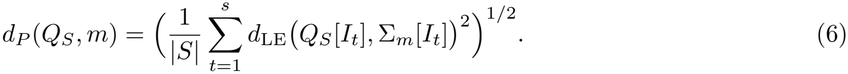

When every nonempty intersection *S* ∩ *B*_*j,b*_ is a single dyadic interval, *d*_*P*_ equals 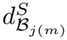; otherwise, *d*_*P*_ omits covariances between distinct pieces of the same block, and Spindle-naive provides the exact reference ranking under 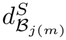 (section B.7).

#### Gene-list queries

For a query given only as a list of genes, Spindle removes genes that are absent from the index and constructs a template covariance matrix over the remaining set *S*:

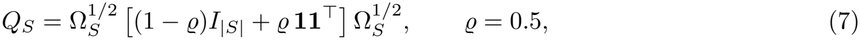

where Ω_*S*_ is diagonal and its entry for each gene is the 90th percentile of that gene’s shrinkage-regularized variance across the index tiles. The template thus uses the index’s gene-wise variance scale and assigns the same pairwise correlation *ϱ* to every pair of queried genes.

## 4 Discussion

We demonstrated that Spindle enables fast retrieval of tissue regions with similar gene–gene covariance patterns without storing the whole covariance matrices from every tile. By grouping tiles into covariance niches and representing each niche with a layered DAG, Spindle reduced storage 113–197-fold compared with storing the whole covariance matrices across eight Xenium datasets. Spindle answered complete queries in 2.8–28 ms on one core (5.1–16 times faster than a whole-matrix scan, and 13–25 times faster on 32 cores), and the returned tiles were close to the exact nearest neighbors, with a median RDE of 11.0% and a median NNR of 81%. For partial queries, Spindle-Partial was 4.0–7.8 times faster than the Spindle-naive search. These results suggest that exploiting the block structure of gene-–gene covariance matrices makes region-level search practical as the number of cells increases.

The query experiments illustrate a large variety of use cases for the index. Within a dataset, the tiles returned by Spindle had cell-type compositions that were significantly closer to that of the query than those of random tiles. Across Visium and Xenium datasets, they had tumor content similar to that of the query, and gene-list queries retrieved tiles with higher expression scores for all 24 curated signatures. A gene-list query requires only the names of the genes, not a covariance matrix measured in the target tissue. Owing to the low disk footprint of the Spindle index, we envision an atlas-scale distribution of pre-indexed ST datasets. Such an indexed atlas will enable the search for arbitrary gene–gene covariance patterns across many tissues, disease conditions and patients.

While Spindle is, to our knowledge, the first of its kind, we identify a few drawbacks in our current implementation. First, Spindle ranks candidate tiles by their distances to node representatives rather than to their own block logarithms; a broader search budget reduced the RDE of complete queries to 5.4–11.1%, but did not eliminate this approximation error. Second, the block distance ignores covariances between blocks and includes the gene variances on each block’s diagonal, so it may partly reflect cell-type composition; distances that normalize or separately weight variances and correlations could better distinguish the two. Third, queries transferred across tissues or platforms required an adjustment: without the global adjustment, the correlation between the tumor fraction of Xenium queries and that of the retrieved Visium tiles decreased from 0.73 to 0.07. Accounting for gene- and platform-specific effects could improve retrieval robustness. Finally, our evaluation covers eight Xenium datasets with 313–480 genes and one paired Xenium–Visium dataset; the performance of Spindle on other platforms and on whole-transcriptome assays should be investigated as a part of large-scale benchmarking.

The block decomposition provides a scalable path to larger gene panels by reducing storage growth from quadratic to linear in the number of genes. It also enables parallel computation and more efficient indexed search, with the greatest benefits expected for datasets with larger gene panels. Furthermore, a shared gene ordering and block structure could enable a unified index across datasets, to which new tiles could be added without rebuilding the index. Overall, Spindle establishes covariance-based indexing as a practical approach to region-level search in ST collections, and these extensions could carry it to larger panels, additional platforms and multi-dataset collections.

## Acknowledgements

Hirak Sarkar and Souvik Ghosh are partially supported by IRG-22-145-62 and the VICC GI SPORE (P50CA236733). This work used TAMU ACES, through allocation CIS250912: Learned Indices for Biomedical Spatial Data from the Advanced Cyberinfrastructure Coordination Ecosystem: Services & Support (ACCESS) program, which is supported by U.S. National Science Foundation grants #2138259, #2138286, #2138307, #2137603, and #2138296.

## Supplementary Material

Spindle: Scalable Indexing of Spatial-Omics Data for Fast Querying

### A Supplementary results

#### A.1 Block structure of the covariance matrices

To examine how much Spindle leaves out by comparing tiles block by block, we convert each tile’s shrunk covariance matrix to a correlation matrix, order its genes as in the tile’s niche, and compute the mean absolute correlation over the gene pairs within blocks and over the pairs between blocks (Supplementary Fig. S1). Pairs involving a gene without counts in the tile are excluded, since their correlation is undefined. In the breast-cancer dataset, the mean absolute correlation is 0.18 within blocks and 0.06 between blocks, and it is higher within blocks for every index tile. The same holds for every tile of the other seven datasets, with the smallest difference for the 5k lymph-node dataset (0.12 against 0.085).

**Fig. S1:**
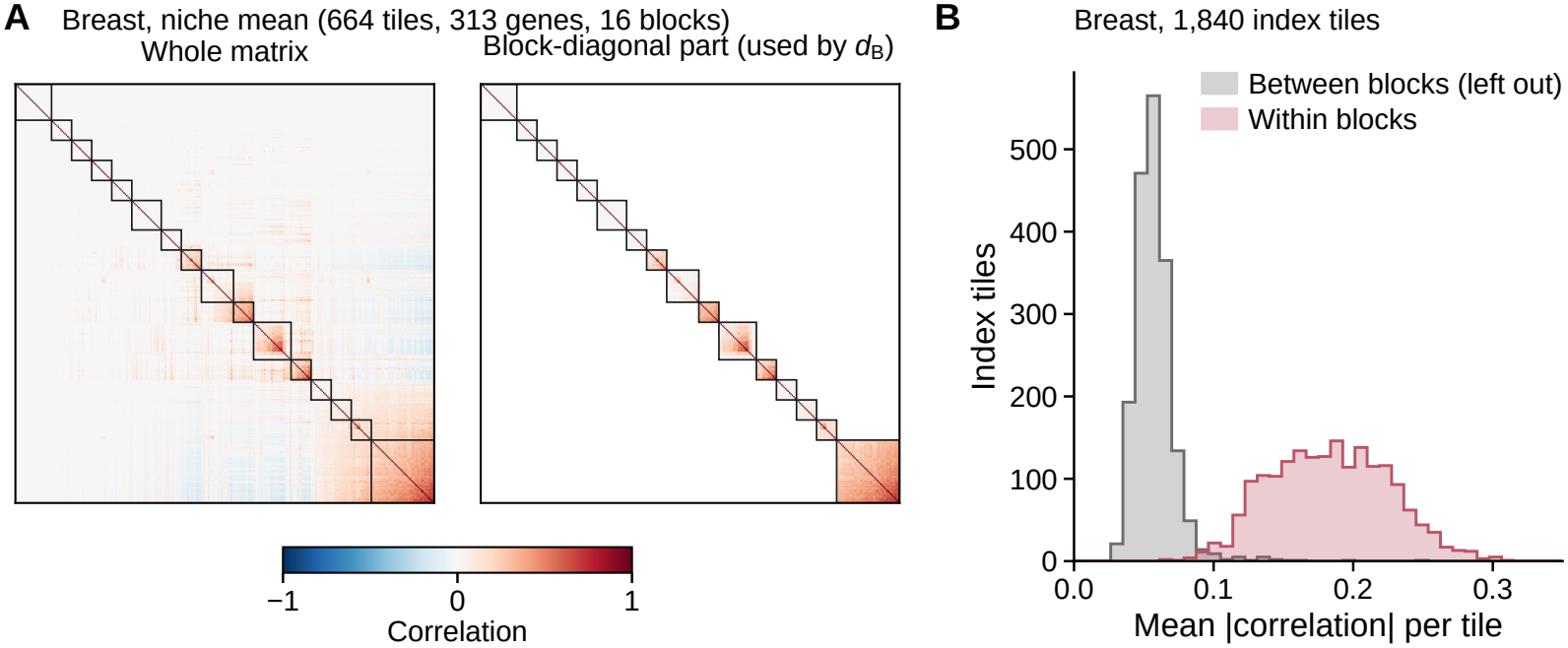
Block structure of the breast-cancer covariance matrices. **A** Mean correlation matrix of the largest covariance niche (664 index tiles, 313 genes, 16 blocks) in the niche’s gene order: whole (left) and its block-diagonal part, which *d*_*ℬ*_ uses (right). **B** Mean absolute correlation of each of the 1,840 index tiles over its within-block gene pairs (red) and between-block gene pairs (gray), each tile in the gene order of its niche; pairs with a gene without counts in the tile are excluded.

To check that leaving out the between-block covariances preserves the order of the tiles, which makes the exact ranking of Spindle-naive under *d*_*ℬ*_ a suitable reference for the accuracy of Spindle, we compare *d*_*ℬ*_ with the whole-matrix log-Euclidean distance on the same scale, 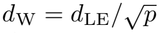, for 1,000 random pairs of distinct index tiles in each of the three largest breast-cancer niches (Supplementary Fig. S2). The two distances are almost monotonically related, with Spearman correlations of 0.942–0.955 and Pearson correlations of 0.969– 0.985. *d*_*ℬ*_ is smaller than *d*_W_ for 98.0–99.7% of the pairs, by a median of 0.19–0.30, and this gap does not grow with *d*_W_: omitting the between-block covariances mostly shifts the distances rather than reordering the tiles.

#### A.2 Covariance niches and gene blocks of breast cancer

To see whether the niches and blocks that Spindle uses for indexing also describe the tissue, we build the niches and blocks from all tiles of the breast-cancer dataset (Supplementary Fig. S3). The three niches follow the tissue architecture: niche 1 is enriched for DCIS and myoepithelial cells, niche 2 for invasive tumor cells, and niche 3 for stromal, endothelial and immune cells. The block of a gene depends on the niche; for example, ACTA2 shares a block with myoepithelial genes in niche 1 and with fibroblast genes in niche 2. We call a block co-expressed when the mean absolute correlation of its genes exceeds that of 95% of 1,000 random gene sets of the same size; per niche, 6–8 of the 15–17 blocks are co-expressed, and 3–6 are enriched for a Hallmark [41] or GO Biological Process [42, 43] term (hypergeometric test on the Enrichr gene-set libraries [44], Benjamini–Hochberg false discovery rate [45] below 0.05).

**Fig. S2:**
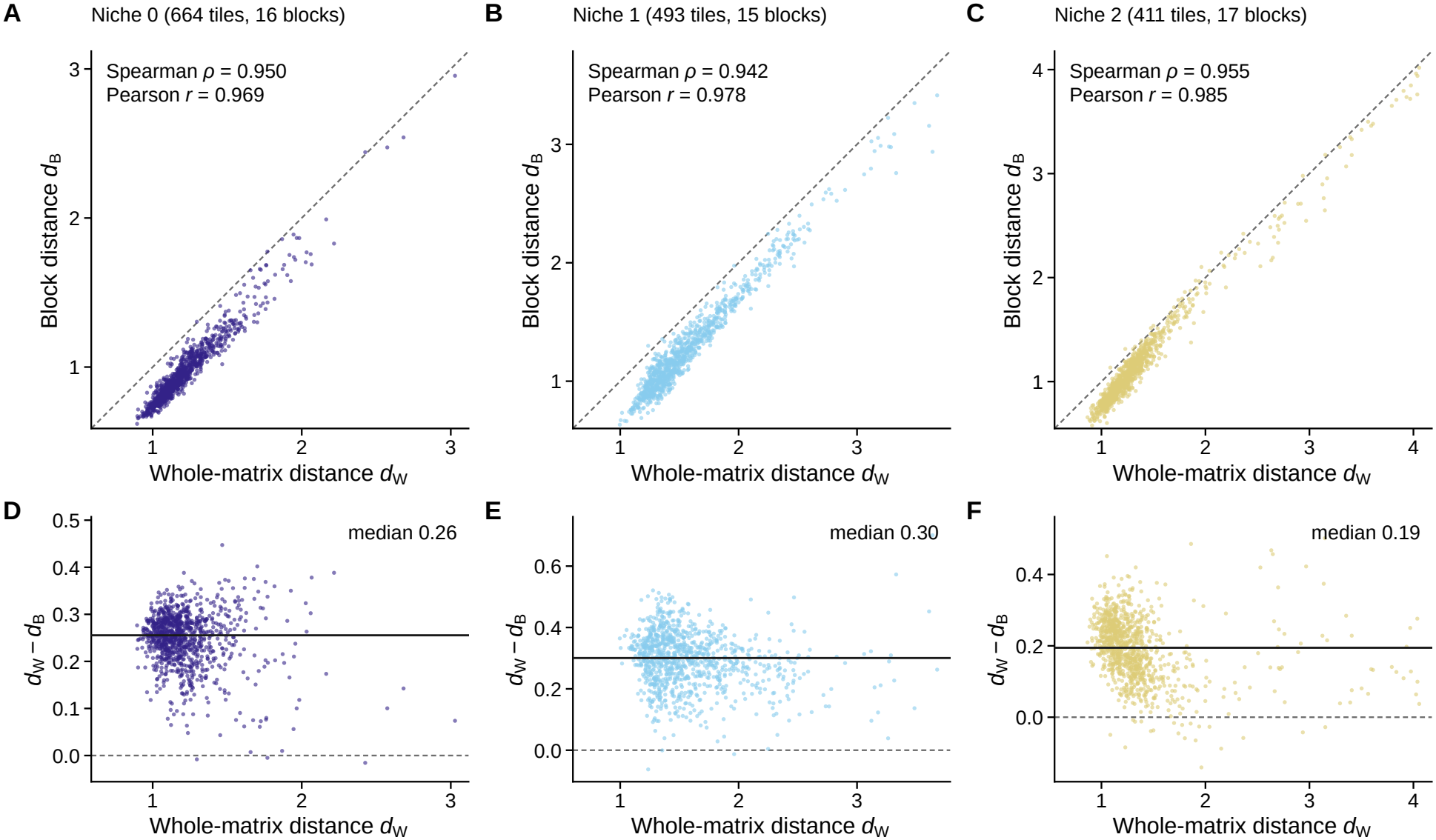
Block distance against whole-matrix distance in breast cancer. Each point is one of 1,000 random pairs of distinct index tiles of a covariance niche (split with seed 0); columns, the three largest niches, with niche 0 as in Supplementary Fig. S1A. **A–C** Block distance *d* in the niche’s gene order and blocks against the whole-matrix log-Euclidean distance 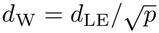; dashed line, *d*_*ℬ*_ = *d*_W_. Insets give the Spearman and Pearson correlations. **D–F** Difference *d*_W_ − *d*_*ℬ*_ against *d*_W_; solid line, median; dashed line, zero.

#### A.3 Datasets and index construction

Supplementary Table S1 lists the size of each dataset, the covariance niches and blocks of its index, the storage of each index, and the time of each construction step. The covariance niches and block partition (layout) take most of the construction time.

#### A.4 Index cost against the number of cells and genes

To measure how the cost grows with the size of a dataset, we build the indexes on a Xenium tonsil dataset [46] (1.35 million cells, 377 genes) subsampled from 1,000 cells to all cells, and on the 5k lymph-node dataset restricted to its 250–1,500 most variable genes (Supplementary Fig. S4). Every index grows in proportion to the number of index tiles, the Spindle index by about 2 kB per tile. With the number of genes, the whole covariance matrices grow quadratically, from 0.87 to 31 GB, whereas the Spindle index grows from 10 to 42 MB; with many genes, building Spindle-naive dominates the construction time.

#### A.5 Computing environment and query time

All indexes were built and all queries timed on one compute node (Supplementary Table S2). Supplementary Tables S3 and S4 list, for complete and partial queries, the time of query preprocessing (the logarithms of the query) and the time of each search. Query preprocessing runs once per query and takes longer than the Spindle search itself.

**Fig. S3:**
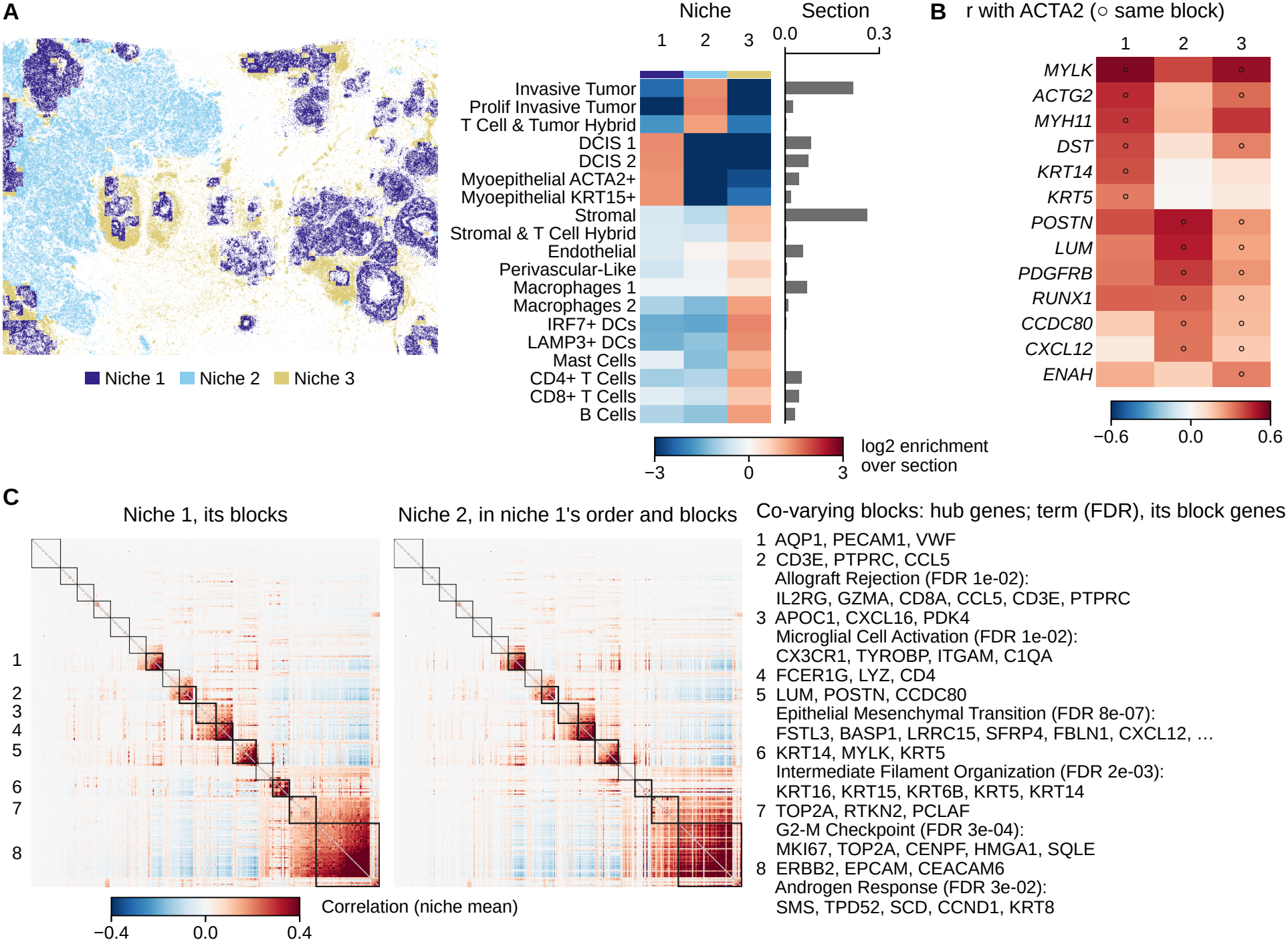
Covariance niches and gene blocks of the breast-cancer dataset. **A** Left, every cell colored by the covariance niche of its tile (light gray, cells of discarded low-density tiles). Middle, the log2 enrichment of each annotated cell type in the cells of each niche relative to the whole dataset. Right, the fraction of each cell type in the dataset. **B** The correlation of ACTA2 with its block partners in each niche, in the mean correlation matrix of the niche; circles, genes in the block of ACTA2 in that niche. **C** Left, the mean correlation matrix of niche 1 in its gene order, with its blocks outlined (thick, co-expressed blocks, numbered). Middle, the mean correlation matrix of niche 2 in the gene order and blocks of niche 1. Right, the co-expressed blocks of niche 1: hub genes (the three genes with the largest summed absolute correlation to the rest of the block), and for enriched blocks the key term, its false discovery rate and its block genes. All panels use niches and blocks built from all tiles of the dataset.

We also timed complete queries on 1–64 physical cores of the same node, using one hardware thread per core (Supplementary Fig. S5). For Spindle, query preprocessing (the block logarithms of the query) is distributed over the cores, the niches of the search are processed on separate cores, and the results are combined into the same ranking as on one core. For the whole-matrix scan, query preprocessing (the logarithm of the whole covariance matrix) uses several threads of the linear-algebra library, and the scan divides the index tiles among the cores. Every parallel step returns the same values as its single-core counterpart. Small dense eigendecompositions do not benefit consistently from additional threads; we therefore report each step at its fastest core count up to the plotted number (Supplementary Fig. S5). Query preprocessing of the whole-matrix scan took 4.9–14 ms at every number of cores and became its floor, whereas that of Spindle decreased to 0.33–0.99 ms on 32 cores, and its time for query preprocessing and search leveled off at 0.46–2.0 ms.

#### A.6 Retrieval accuracy

For a typical breast-cancer query, the tiles returned by Spindle lie in the lower tail of the distance distribution, for complete and partial queries alike (Supplementary Fig. S6A,C). Increasing the search-budget breadth *g* from its default of 0.25 lowers the RDE of complete queries on every dataset, to 5.4–11.1% at the largest value tested (Supplementary Fig. S6B). For partial queries, the RDE of Spindle-Partial increases from 8.6% with 4 genes to 18.9% with 32, whereas padding and imputation remain far from the exact answer at every query size (Supplementary Fig. S6D,E).

**Table S1:** Datasets and indexes. Top, the size of each dataset, the covariance niches and gene blocks of its index (range over the five splits), and the storage of the whole covariance matrices of the index tiles, Spindle-naive, Spindle, and their indexes for partial queries (Spindle-naive (P) and Spindle-Partial). Bottom, construction time per step of building all four indexes: reading the dataset, tiles and their covariance matrices, layout (covariance niches, gene orders and blocks), and each index; the total excludes reading. Means over the five splits.

|  |  |  | Tiles |  |  |  |  | Storage (MB) |  |  |  |  |
|---|---|---|---|---|---|---|---|---|---|---|---|---|
| Dataset | Cells | Genes | Tiles | Index | Held out | Niches | Blocks | Whole | Spindle-naive | Spindle | Spindle-naive (P) | Spindle-Partial |
| Skin | 87,499 | 382 | 932 | 839 | 93 | 3–4 | 58–82 | 245.5 | 15.9 | 1.8 | 31.8 | 2.2 |
| Kidney | 97,560 | 377 | 783 | 705 | 78 | 2–3 | 40–60 | 200.9 | 11.3 | 1.3 | 22.5 | 1.6 |
| Breast | 159,226 | 313 | 2,044 | 1,840 | 204 | 3–4 | 45–67 | 361.7 | 28.5 | 3.2 | 57.0 | 3.9 |
| Lung | 162,254 | 377 | 1,705 | 1,535 | 170 | 3–4 | 57–77 | 437.5 | 28.8 | 3.2 | 57.5 | 3.9 |
| Pancreas | 190,965 | 474 | 2,290 | 2,061 | 229 | 4 | 108–111 | 928.1 | 39.3 | 4.7 | 78.5 | 5.7 |
| Lymph node | 377,985 | 377 | 2,615 | 2,354 | 261 | 5–7 | 103–146 | 670.9 | 41.2 | 4.5 | 82.4 | 5.7 |
| Lymph node 5k | 708,983 | 400 | 7,746 | 6,972 | 774 | 15–17 | 302–345 | 2,236.6 | 136.8 | 15.7 | 273.6 | 18.8 |
| Brain | 816,769 | 480 | 10,620 | 9,558 | 1,062 | 16–18 | 418–480 | 4,413.5 | 201.6 | 23.3 | 403.1 | 28.5 |
|  | Build time (s) |  |  |  |  |  |  |  |  |  |  |  |
| Dataset | Read | Tiles + covs | Layout | Spindle-naive | Spindle | Spindle-naive (P) | Spindle-Partial | Total |  |  |  |  |
| Skin | 3.8 | 0.6 | 19.6 | 1.0 | 0.7 | 0.6 | 4.8 | 27.3 |  |  |  |  |
| Kidney | 4.0 | 0.5 | 18.2 | 0.8 | 0.6 | 0.4 | 3.4 | 23.8 |  |  |  |  |
| Breast | 4.7 | 1.2 | 23.9 | 1.7 | 1.4 | 1.0 | 7.1 | 36.4 |  |  |  |  |
| Lung | 4.0 | 1.0 | 25.0 | 1.8 | 1.4 | 1.0 | 6.7 | 36.9 |  |  |  |  |
| Pancreas | 4.6 | 3.2 | 42.0 | 3.2 | 2.1 | 1.4 | 10.8 | 62.6 |  |  |  |  |
| Lymph node | 4.5 | 1.6 | 33.1 | 2.8 | 2.1 | 1.5 | 11.0 | 52.0 |  |  |  |  |
| Lymph node 5k | 7.4 | 7.8 | 102.4 | 10.2 | 6.9 | 5.6 | 35.1 | 168.0 |  |  |  |  |
| Brain | 7.4 | 8.3 | 169.1 | 14.5 | 10.5 | 6.9 | 50.4 | 259.7 |  |  |  |  |

**Fig. S4:**
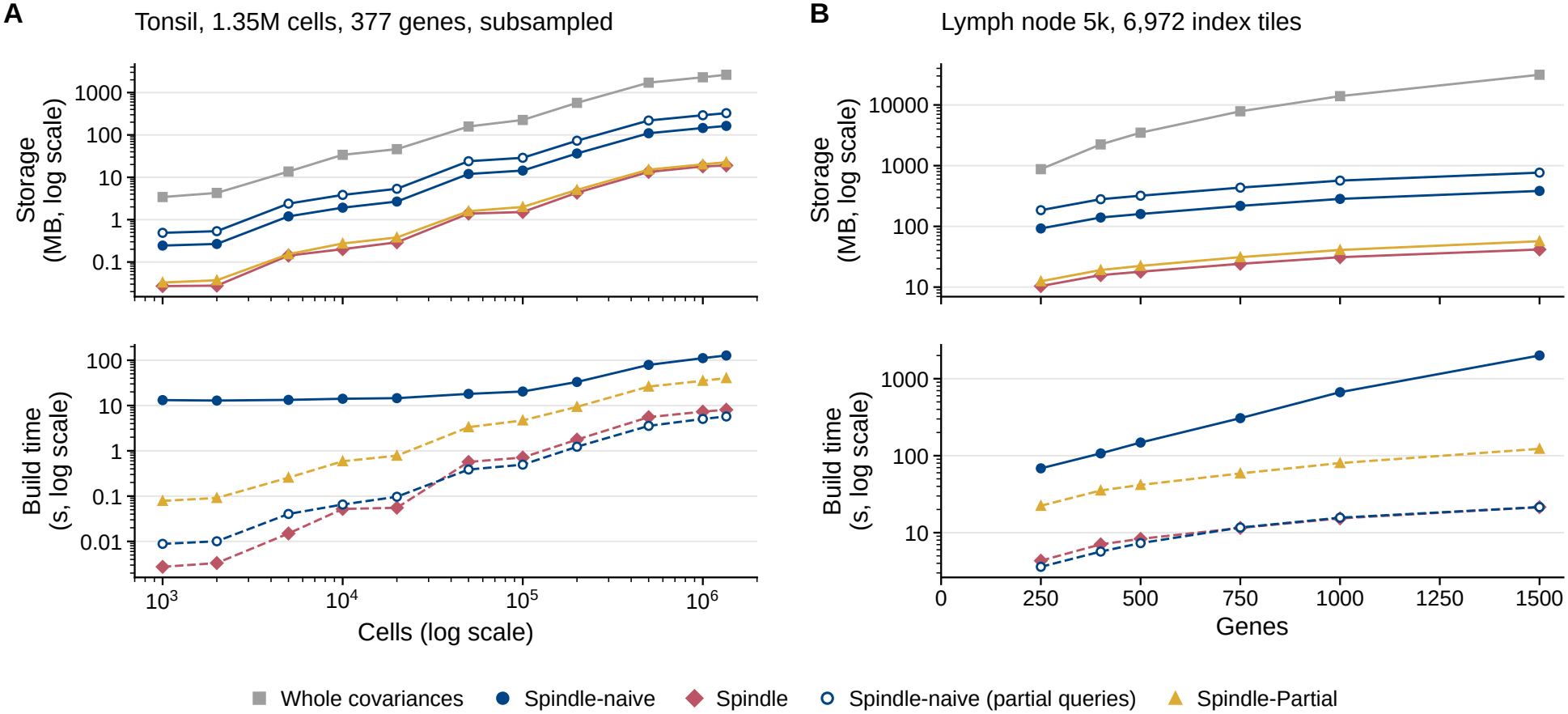
Index cost against the number of cells and genes. **A** A Xenium tonsil dataset (1.35 million cells, 377 genes) subsampled to 1,000–1.35 million cells: storage of each index (top) and construction time (bottom; Spindle-naive including tiles, covariance matrices and layout, solid; the time each other index adds, dashed). **B** The same for the 5k lymph-node dataset (6,972 index tiles) restricted to its 250–1,500 most variable genes.

**Table S2:**
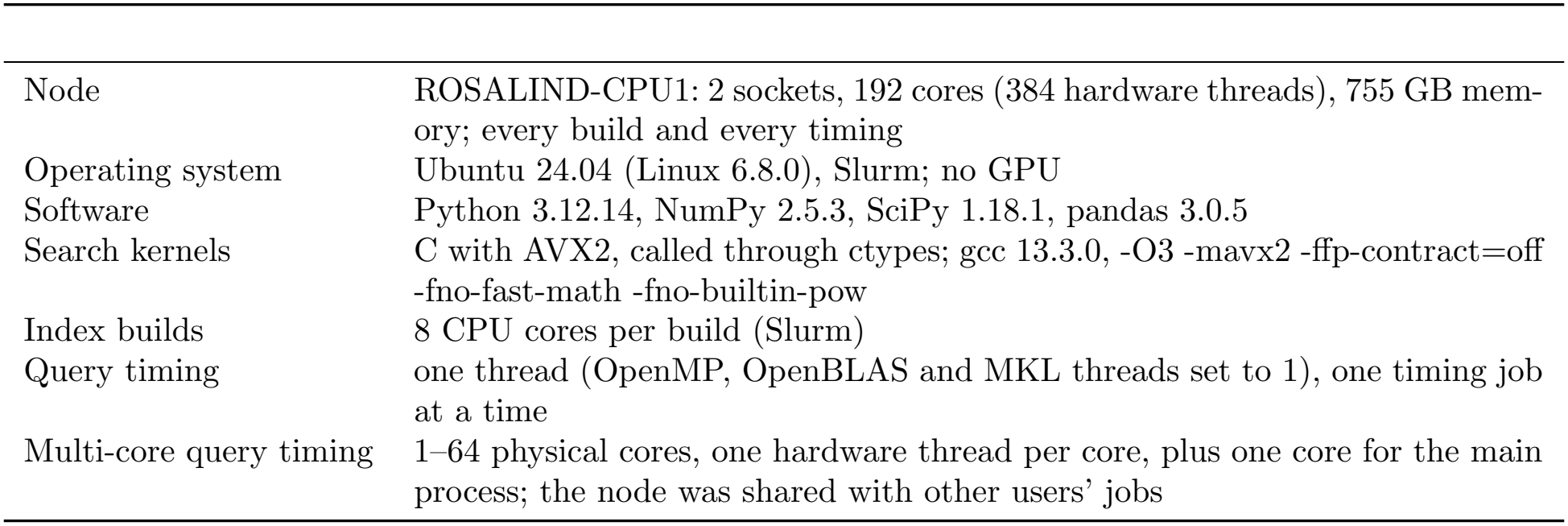
Computing environment.

**Table S3:** Time of a complete query. Read, time to read the dataset (s). Per query (ms): query preprocessing of the whole-matrix scan (the logarithm of the whole covariance matrix), the scan over the whole-matrix logarithms of the index tiles, and their total; query preprocessing of Spindle-naive and Spindle (the block logarithms of the query in every niche, shared by both); the two searches; and the total for Spindle (query preprocessing + search). Mean and standard deviation across the five splits of the median over the first 200 held-out queries (all, if fewer); one core.

|  |  | Whole matrix (ms) |  |  | Block representation (ms) |  |  |  |
|---|---|---|---|---|---|---|---|---|
| Dataset | Read (s) | Logarithm | Scan | Total | Block logarithms | Spindle-naive | Spindle | Spindle total |
| Skin | 3.8 ± 0.4 | 8.9 ± 0.4 | 16.1 ± 0.1 | 25.1 | 4.7 ± 0.6 | 1.12 ± 0.06 | 0.25 ± 0.00 | 4.9 |
| Kidney | 4.0 ± 0.3 | 8.4 ± 0.1 | 14.5 ± 0.1 | 22.8 | 2.6 ± 0.6 | 0.77 ± 0.08 | 0.17 ± 0.02 | 2.8 |
| Breast | 4.7 ± 0.5 | 5.6 ± 0.0 | 23.9 ± 0.2 | 29.4 | 3.8 ± 0.4 | 1.71 ± 0.10 | 0.42 ± 0.02 | 4.2 |
| Lung | 4.0 ± 0.2 | 8.5 ± 0.0 | 30.1 ± 0.2 | 38.5 | 4.8 ± 0.5 | 2.03 ± 0.07 | 0.51 ± 0.02 | 5.3 |
| Pancreas | 4.6 ± 0.4 | 14.4 ± 0.1 | 73.7 ± 0.1 | 88.0 | 6.0 ± 0.0 | 2.87 ± 0.03 | 0.55 ± 0.01 | 6.6 |
| Lymph node | 4.5 ± 0.7 | 8.8 ± 0.0 | 42.3 ± 0.2 | 51.1 | 7.5 ± 0.9 | 2.70 ± 0.16 | 0.67 ± 0.02 | 8.2 |
| Lymph node 5k | 7.4 ± 3.0 | 11.3 ± 0.1 | 114.5 ± 0.1 | 125.8 | 22.1 ± 1.3 | 7.64 ± 0.21 | 2.38 ± 0.07 | 24.5 |
| Brain | 7.4 ± 0.5 | 14.7 ± 0.1 | 441.3 ± 1.6 | 456.0 | 25.5 ± 1.2 | 15.10 ± 0.37 | 2.91 ± 0.08 | 28.4 |

**Table S4:** Time of a partial query. Per query (ms): query preprocessing (the query logarithms) and the scan of Spindle-naive; query preprocessing (the logarithms of the query’s dyadic intervals), the search and their total for Spindle-Partial; and the total time of padding and imputation (completion, logarithms and Spindle-naive search). Queries over 4, 8, 16 and 32 genes pooled; mean and standard deviation across the five splits of the median over the queries of the first 200 held-out tiles (all, if fewer); one core.

|  | Spindle-naive (ms) |  | Spindle-Partial (ms) |  |  | Completion (ms) |  |
|---|---|---|---|---|---|---|---|
| Dataset | Block logarithms | Scan | Piece logarithms | Search | Total | Padding | Imputation |
| Skin | 0.52 ± 0.07 | 4.1 ± 0.2 | 1.03 ± 0.14 | 0.101 ± 0.002 | 1.12 | 4.1 ± 0.4 | 7.7 ± 0.7 |
| Kidney | 0.25 ± 0.05 | 3.4 ± 0.3 | 0.50 ± 0.12 | 0.080 ± 0.008 | 0.58 | 2.5 ± 0.4 | 4.8 ± 0.7 |
| Breast | 0.50 ± 0.07 | 8.9 ± 0.7 | 1.04 ± 0.12 | 0.159 ± 0.004 | 1.20 | 4.1 ± 0.2 | 7.0 ± 0.5 |
| Lung | 0.53 ± 0.08 | 7.5 ± 0.5 | 1.04 ± 0.15 | 0.151 ± 0.005 | 1.19 | 5.2 ± 0.4 | 8.7 ± 0.8 |
| Pancreas | 0.62 ± 0.04 | 6.0 ± 0.6 | 1.07 ± 0.04 | 0.158 ± 0.009 | 1.21 | 7.1 ± 0.1 | 11.2 ± 0.1 |
| Lymph node | 0.90 ± 0.13 | 10.8 ± 0.7 | 1.69 ± 0.25 | 0.214 ± 0.006 | 1.90 | 7.8 ± 0.6 | 12.5 ± 1.1 |
| Lymph node 5k | 2.68 ± 0.17 | 30.1 ± 2.7 | 4.60 ± 0.27 | 0.541 ± 0.008 | 5.14 | 20.7 ± 0.9 | 34.0 ± 1.6 |
| Brain | 2.69 ± 0.14 | 25.6 ± 2.6 | 4.83 ± 0.22 | 0.454 ± 0.015 | 5.29 | 33.0 ± 0.8 | 44.9 ± 1.4 |

**Fig. S5:**
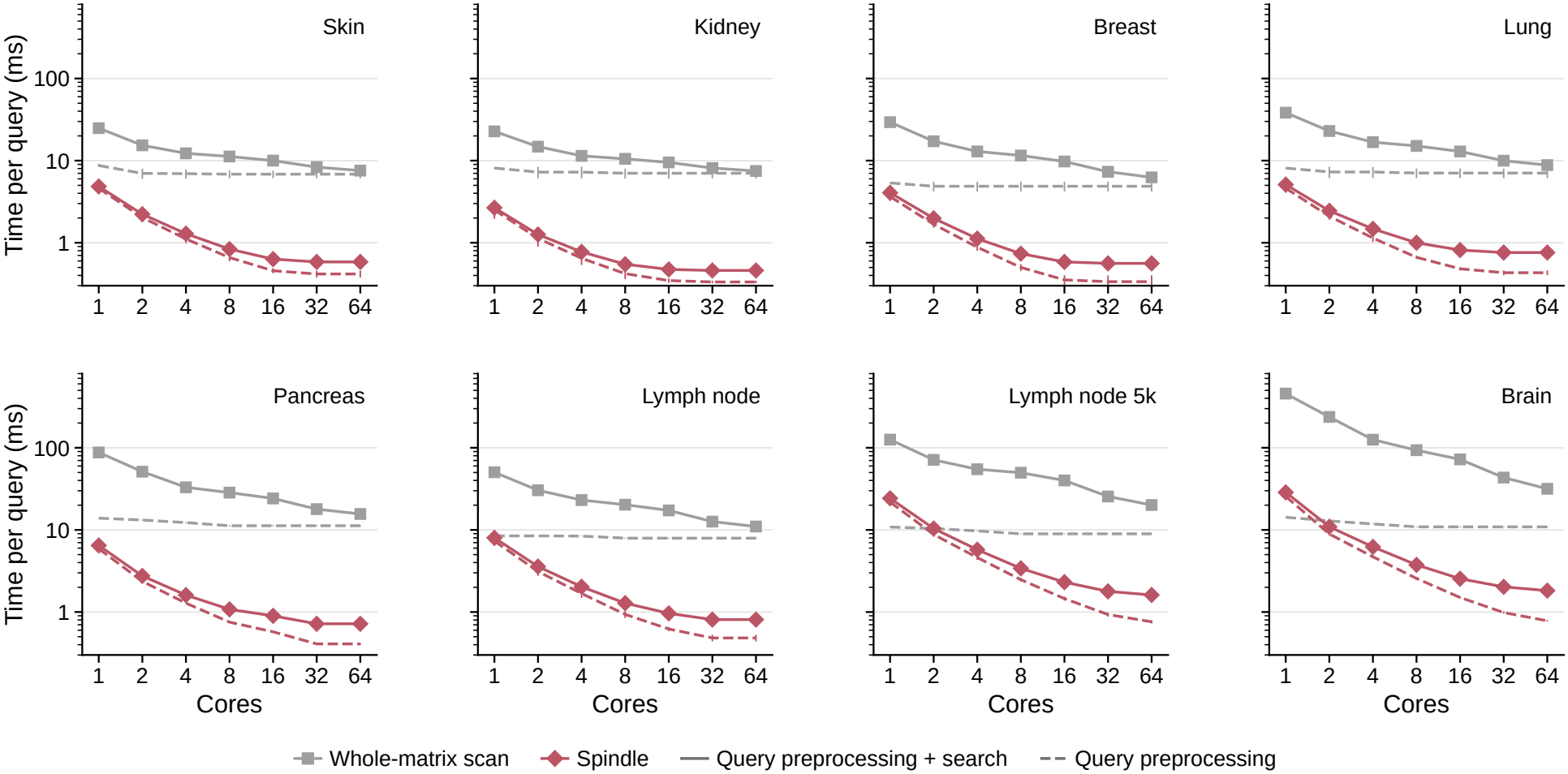
Time of a complete query on several cores. Time per query (log scale) of the whole-matrix scan and Spindle against the number of physical cores, for each dataset: query preprocessing + search (solid) and query preprocessing alone (dashed; the logarithm of the whole covariance matrix for the scan, the block logarithms of the query for Spindle). Each step is shown at its fastest number of cores up to the plotted number. Mean and standard deviation across five random splits of the median over the first 200 held-out queries; the node was shared with other jobs.

#### A.7 Platform shift and tissue agreement

##### Global adjustment

Before a query from one platform is searched against an index of the other, the logarithm *L* of the query’s whole covariance matrix is replaced by the affine map

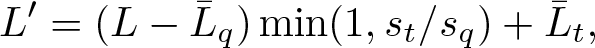

where 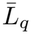 and 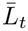 are the entry-wise means of the logarithms of the query and index tiles, and *s*_*q*_ and *s*_*t*_ are their standard deviations over all entries; the covariance matrix exp *L*′ is then searched. The adjustment uses no cell-type annotations.

##### Platform shift

In the first two principal components of the covariance logarithms, fitted on the Xenium tiles, the Visium tiles are offset from the Xenium tiles, and the global adjustment moves them together (Supplementary Fig. S7A). After the adjustment, the tumor fraction of a query and the mean tumor fraction of its first 10 retrieved tiles on the other platform increase together for queries from either platform (Supplementary Fig. S7B). Without adjustment, the tumor fractions of the queries and of their retrieved tiles do not agree, and a separate adjustment in every block of every niche gives lower agreement than the global adjustment (Supplementary Fig. S7C).

**Fig. S6:**
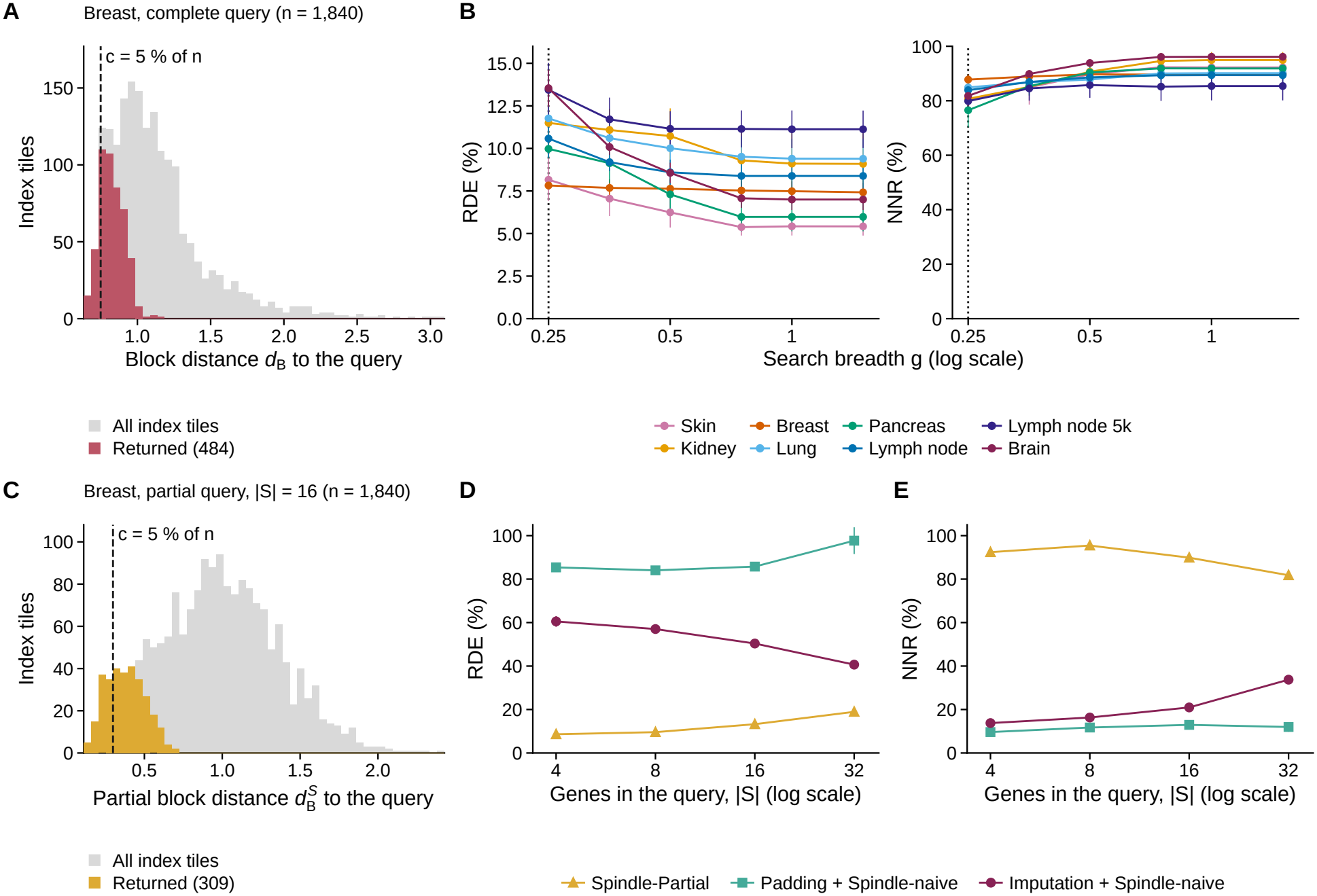
Retrieval accuracy. **A** A typical breast-cancer complete query: *d*_*B*_ from the query to every index tile (gray) and to the tiles returned by Spindle (red); dashed line, the distance of the *c*-th nearest index tile at *c* = 5% of the index size *n*. **B** RDE (left) and NNR (right) of Spindle against the search-budget breadth *g* (log scale; dotted line, the default *g* = 0.25) for each dataset. **C** As **A**, for a partial query over 16 genes and the partial block distance 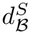. **D, E** RDE and NNR against the number of query genes |*S*| (log scale) for Spindle-Partial and for padding and imputation followed by a Spindle-naive search; per split, the median over the eight datasets. **B, D, E** Mean and standard deviation across five random splits.

#### A.8 Partial and gene-list queries on every dataset

##### Cross-dataset partial queries

Each query of Fig. 4A takes its gene–gene correlations from one breast-cancer tile and its gene variances from the index it searches. Among the breast-cancer index tiles with at least 80 cells and at least 0.5 counts of the query genes *S* per cell, the exemplar is the tile with the largest mean correlation between distinct genes of *S* across its cells, provided that no cell holds more than 15% of the counts of *S* and that removing its three cells with the most counts keeps at least 70% of that correlation. Let *R*_*S*_ be the correlation matrix of the exemplar’s shrunk covariance matrix restricted to *S*, and *q*_*g*_ the fraction of breast-cancer index tiles whose shrunk variance of gene *g* does not exceed that of the exemplar. In a target dataset, the query is *Q*_*S*_ = diag(*σ*) *R*_*S*_ diag(*σ*), where 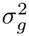 is the *q*_*g*_-quantile of the shrunk variance of gene *g* across the index tiles of that dataset, so that every gene keeps the rank of its exemplar variance. Closeness is 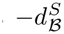 from Spindle-naive, and the expression score of a tile is the mean of the z-scored mean log-normalized expression of the genes of *S*. Supplementary Table S5 gives the correlation between closeness and expression score on every dataset whose index contains all the query genes: it is positive for the myeloid genes on every dataset, but negative for the cytotoxic T-cell genes on the kidney, pancreatic-cancer and brain-cancer datasets.

**Fig. S7:**
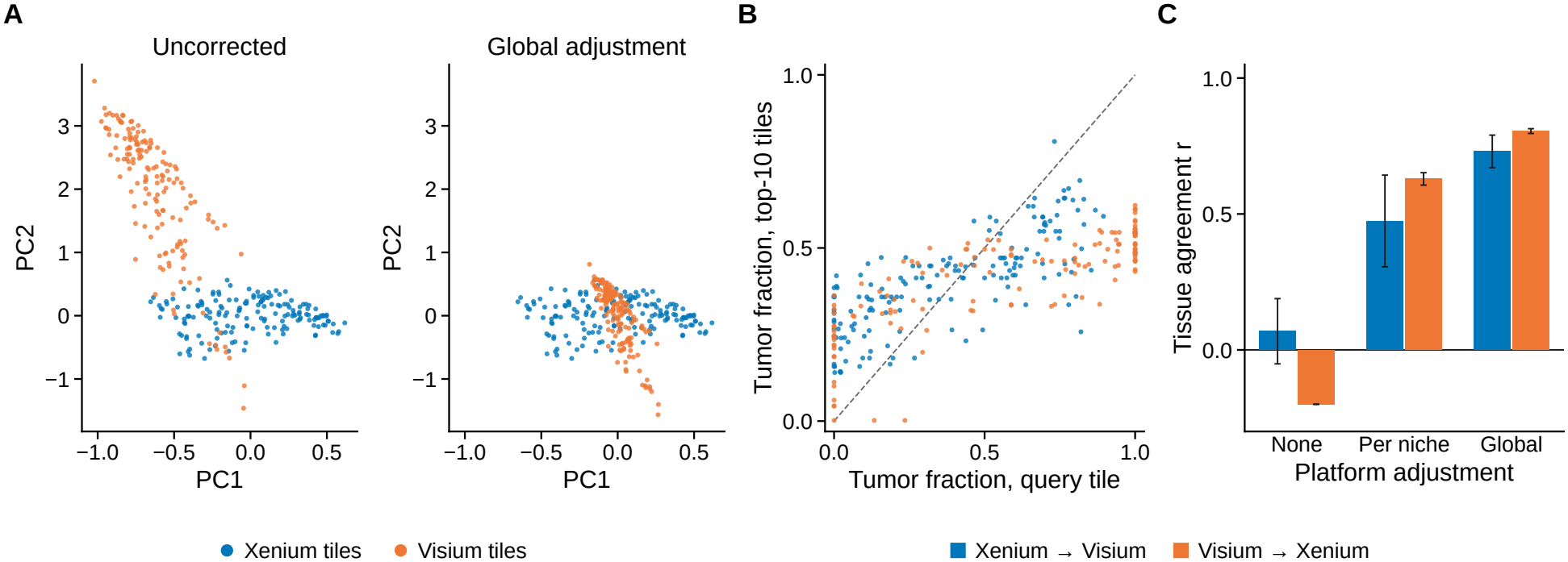
Platform shift and tissue agreement between Xenium and Visium. **A** The first two principal components of the logarithms of the whole covariance matrices of the Xenium tiles (blue) and the Visium tiles (orange), fitted on the Xenium tiles, before (left, Uncorrected) and after (right) the global adjustment of the Visium tiles. **B** The tumor fraction of every query tile against the mean tumor fraction of the first 10 tiles returned by Spindle on the other platform, after the global adjustment; Xenium queries searched against the Visium tiles (blue) and Visium queries against the Xenium tiles (orange); queries for which Spindle returned no tile are left out. Dashed line, the diagonal. **C** Tissue agreement, the Pearson correlation of **B**, without adjustment (None), with an adjustment per block of every niche (Per niche), and with the global adjustment (Global). Queries for which Spindle returned no tile are left out. Every tile is indexed; mean and standard deviation over five random seeds of the niche construction.

##### Partial and gene-list queries

Supplementary Table S6 repeats the within-dataset partial-query comparison of Fig. 4B on every dataset. Supplementary Table S7 lists the 24 curated signatures, the genes of each that are present in the index, and the enrichment of the tiles retrieved by the corresponding gene-list queries.

**Table S7:** Curated tissue signatures searched as gene-list queries. For each signature: its class, the genes present in the index, Cliff’s *δ* between the expression scores of the retrieved tiles and those of the remaining index tiles (mean and standard deviation across five random splits), and the mean number of tiles retrieved (at most 10; fewer when the distance budget returns fewer).

| Dataset | Signature | Class | Genes searched | Cliff's δ | Tiles returned |
|---|---|---|---|---|---|
| Skin | Melanocyte Tumor | Tumor / epithelial | TYR, TYRP1, MLANA, PMEL, DCT, MITF, SOX10, S100B | 0.81 ± 0.02 | 8.6 |
| Skin | Keratinocyte Epidermis | Tumor / epithelial | KRT1, KRT5, KRT17, LOR | 0.79 ± 0.02 | 7.0 |
| Skin | Immune Infiltrate | Lymphoid | CD3D, CD3E, CD68, C1QA, ITGAX | 0.88 ± 0.03 | 3.0 |
| Kidney | Tubular Epithelium | Tumor / epithelial | SLC22A8, SLC4A1, AQP2, UMOD, SLC26A3 | 0.65 ± 0.05 | 10.0 |
| Kidney | Injury Immune Infiltrate | Lymphoid | HAVCR2, CD68, CD163, CD3D, CD3E | 0.28 ± 0.12 | 6.4 |
| Breast | Luminal Tumor Core | Tumor / epithelial | ESR1, PGR, ERBB2, FOXA1, GATA3, KRT8, EPCAM, CDH1 | 0.64 ± 0.02 | 10.0 |
| Breast | Macrophage Myeloid | Myeloid | CD68, CD163, MRC1, C1QA, ITGAX | 0.74 ± 0.02 | 7.6 |
| Breast | Basal Myoepithelial | Tumor / epithelial | KRT5, KRT14, ACTA2, MYLK | 0.81 ± 0.01 | 10.0 |
| Breast | Endothelial Vascular | Stroma / vascular | PECAM1, VWF, KDR | 0.68 ± 0.07 | 10.0 |
| Breast | Proliferation Signature | Tumor / epithelial | MKI67, TOP2A | 0.69 ± 0.04 | 10.0 |
| Lung | Alveolar Airway Epithelium | Tumor / epithelial | AGER, FOXJ1, SCGB2A1, KRT7, SOX2 | 0.43 ± 0.02 | 8.8 |
| Lung | Vascular Endothelial | Stroma / vascular | SOX17, SOX18, PECAM1, VWF | 0.71 ± 0.02 | 10.0 |
| Pancreas | Exocrine Ductal | Tumor / epithelial | KRT7, KRT20, CFTR | 0.57 ± 0.03 | 10.0 |
| Pancreas | Endocrine Islet | Tumor / epithelial | INS, GCG, SST, PPY, CHGA | 0.77 ± 0.00 | 10.0 |
| Pancreas | Mast Cell Stroma | Stroma / vascular | CPA3, ACTA2, PDGFRB | 0.87 ± 0.01 | 9.4 |
| Lymph node | B Cell Germinal Center | Lymphoid | MS4A1, CD79A, CD19, MZB1 | 0.51 ± 0.08 | 9.8 |
| Lymph node | T Cell Zone | Lymphoid | CD3D, CD3E, CD4, CD8A | 0.68 ± 0.03 | 8.2 |
| Lymph node | Macrophage DC Niche | Myeloid | CD68, CD163, LAMP3, CD83, CD86, CD300E | 0.70 ± 0.06 | 10.0 |
| Lymph node | Lymphatic Vascular Stroma | Stroma / vascular | LYVE1, PROX1, PDPN, CD34, CCL19 | 0.93 ± 0.02 | 3.0 |
| Lymph node 5k | B Cell Germinal Center | Lymphoid | MS4A1, CD79A, CD19, MZB1 | 0.75 ± 0.05 | 5.8 |
| Lymph node 5k | T Cell Zone | Lymphoid | CD3E, CD4, CD8A | 0.69 ± 0.04 | 6.8 |
| Brain | Glioma Tumor Proliferative | Tumor / epithelial | EGFR, PDGFRA, MKI67 | 0.35 ± 0.07 | 5.0 |
| Brain | Microglia Myeloid | Myeloid | CD68, CD44, PTPRC, MMP9, MMP12 | 0.55 ± 0.03 | 8.6 |
| Brain | TIL Lymphocyte | Lymphoid | CD3D, CD3E, CD37, CD38 | 0.18 ± 0.19 | 6.4 |

**Table S6:** Composition difference of partial queries from held-out tiles. For each dataset and query size |*S*| : the number of queries for which Spindle returned at least one tile (out of all held-out tiles), the median composition difference between a query tile and its first 10 retrieved tiles (Spindle) or 10 random index tiles (Random, averaged over 200 draws), the percentage of queries for which the retrieved tiles are closer, and an upper bound on the one-sided Wilcoxon signed-rank *p*-value. One random split.

|  | \|S\| = 4 |  |  |  |  | \|S\| = 16 |  |  |  |  |
|---|---|---|---|---|---|---|---|---|---|---|
| Dataset | Queries | Spindle | Random | Closer (%) | *p* | Queries | Spindle | Random | Closer (%) | *p* |
| Skin | 93/93 | 0.38 | 0.57 | 84 | < 10<sup>−12</sup> | 93/93 | 0.36 | 0.58 | 88 | < 10<sup>−12</sup> |
| Kidney | 78/78 | 0.33 | 0.35 | 67 | < 10<sup>−3</sup> | 78/78 | 0.31 | 0.35 | 72 | < 10<sup>−4</sup> |
| Breast | 204/204 | 0.38 | 0.53 | 69 | < 10<sup>−9</sup> | 204/204 | 0.43 | 0.53 | 60 | < 10<sup>−5</sup> |
| Lung | 170/170 | 0.39 | 0.44 | 74 | < 10<sup>−11</sup> | 170/170 | 0.38 | 0.45 | 67 | < 10<sup>−9</sup> |
| Pancreas | 229/229 | 0.39 | 0.49 | 72 | < 10<sup>−14</sup> | 229/229 | 0.38 | 0.49 | 75 | < 10<sup>−16</sup> |
| Lymph node | 261/261 | 0.37 | 0.41 | 67 | < 10<sup>−10</sup> | 261/261 | 0.33 | 0.41 | 74 | < 10<sup>−20</sup> |
| Lymph node 5k | 774/774 | 0.37 | 0.40 | 63 | < 10<sup>−19</sup> | 774/774 | 0.33 | 0.40 | 78 | < 10<sup>−65</sup> |
| Brain | 1062/1062 | 0.32 | 0.36 | 72 | < 10<sup>−49</sup> | 1062/1062 | 0.29 | 0.36 | 80 | < 10<sup>−101</sup> |

**Table S5:** Cross-dataset partial queries on every dataset. Spearman’s *ρ* between the closeness of each index tile to the query 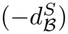 and the mean z-scored expression of the query genes in the tile, for the myeloid and cytotoxic T-cell queries of Fig. 4A, on every dataset whose index contains all the query genes (the 5k lymph-node index, restricted to 400 genes, does not). For breast cancer, the tile from which the query was taken is excluded. One random split.

|  | Spearman's ρ |  |  |
|---|---|---|---|
| Dataset | Myeloid | Cytotoxic T cell | Index tiles |
| Skin | 0.45 | 0.37 | 839 |
| Kidney | 0.37 | -0.08 | 705 |
| Breast | 0.53 | 0.41 | 1,840 |
| Lung | 0.56 | 0.29 | 1,535 |
| Pancreas | 0.28 | -0.15 | 2,061 |
| Lymph node | 0.52 | 0.75 | 2,354 |
| Brain | 0.35 | -0.18 | 9,558 |

### B Supplementary methods

#### B.1 Data, tiles and covariance matrices

##### B.1.1 Data representation and gene selection

We represent a spatial transcriptomics (ST) dataset as *D* = (*Z, X*). Let *N* be the number of cells in the dataset, indexed by *i* ∈ {1, …, *N* }, and let *P* be the number of genes in the panel of the platform. *Z* ∈ ℝ*N ×*2 is the matrix of spatial coordinates, whose *i*-th row *z*_*i*_ ∈ ℝ2 holds the two coordinates of cell *i* on the tissue section. 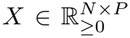 is the count matrix, whose *i*-th row 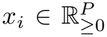 holds the raw transcript counts of the *P* genes in cell *i*, and *x*_*ig*_ denotes the count of gene *g* in cell *i*. Spindle uses raw counts obtained from the experiment.

For a typical Xenium experiment we use all ∼ 400 genes, whereas for the Xenium 5k panel (*P* = 4,624), Spindle uses the *p* = 400 genes with the largest variance of their counts across cells, where the variance of gene *g* is

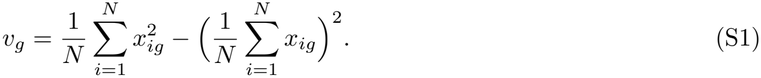

##### B.1.2 Spatial tiles

We divide the ST dataset into axis-aligned rectangles *R* = [*x*_0_, *x*_1_] × [*y*_0_, *y*_1_], termed *tiles*, through a recursive process called *tiling*. We use a capacity-constrained quadtree [24, 25] that starts from the bounding box containing the entire dataset. Each rectangle containing more than *ℓ*_max_ = 200 cells is divided into four quadrants by splitting at the midpoint of each axis. Empty quadrants are discarded, and the remaining quadrants are recursively subdivided until each contains at most *ℓ*_max_ cells. If subdivision cannot proceed or reaches the maximum depth of 40 levels, the cells are partitioned into groups of at most *ℓ*_max_ cells that share the same rectangle.

Tiles of very low cell density occur at the edges of the tissue and in debris, and their covariance matrices carry little co-variation. Spindle removes them with a robust outlier rule. For a tile *T* with box *R*, let |*T* | be its number of cells, |*R*| the area of *R*, and

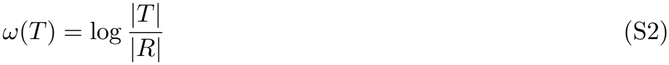

its log cell density. Let med be the median of *ω* over all tiles and MAD the median of |*ω*(*T* ) − med | over all tiles. A tile is removed when *ω*(*T* ) *<* med −6 × 1.4826 × MAD, where the factor 1.4826 makes the MAD a consistent estimate of the standard deviation under a normal distribution; when MAD = 0, no tile is removed. The cells of removed tiles take no part in any later step. We denote the remaining tiles by *T*_1_, …, *T*_*M*_, where each *T*_*m*_ ⊆ {1, …, *N* } is the set of cells of tile *m* and *M* is the number of tiles.

##### B.1.3 Tile covariance matrices

For each tile *T*_*m*_, let 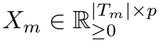 be the matrix whose rows are the count vectors of the cells of *T*_*m*_, restricted to the genes of *G* in the reference order. Let **1** be the all-ones vector of length 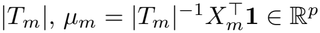 the vector of mean counts of the genes over the cells of the tile, and *I*_*p*_ the *p* × *p* identity matrix. The covariance matrix of tile *m* is

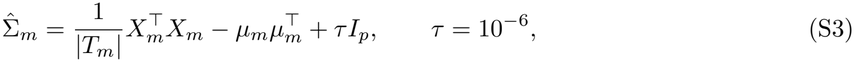

a *p* × *p* matrix whose entry (*g, g*′) is the covariance of the counts of genes *g* and *g*′ across the cells of the tile, normalized by the number of cells, plus a small ridge *τ* on the diagonal.

###### Covariance matrix transformation

A tile contains at most *ℓ*_max_ = 200 cells, whereas *p* is several hundred, so the empirical part of 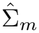 has rank smaller than *p*, and many genes have zero or near-zero counts within a tile. The smallest eigenvalues of 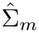 are therefore close to *τ* and would dominate any distance computed on matrix logarithms. Spindle regularizes every tile covariance by shrinking it towards a multiple of the identity [47],

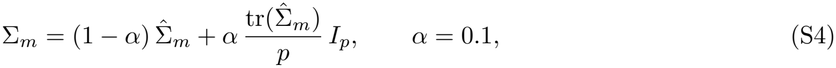

where tr(·) is the trace of a square matrix, so that 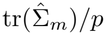 is the average gene variance of the tile. Let 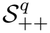 denote the set of *q* × *q* symmetric positive definite (SPD) matrices. Because 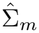 is symmetric positive semi-definite, every eigenvalue of Σ_*m*_ is at least 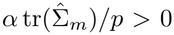, and hence 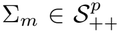. In contrast to the Ledoit–Wolf estimator [47], the weight *α* is fixed rather than estimated per matrix, so that every tile, and every query, is shrunk by the same amount.

Shrinkage is applied once, when each covariance matrix is computed. All subsequent steps use the resulting matrices Σ_*m*_ without further shrinkage. Query covariances computed from tissue regions use the same procedure (Eqs. (S3) and (S4)). We refer to Σ_*m*_ as the covariance of tile *m* and denote the collection of tile covariances by 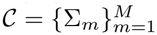.

###### Evaluation

To evaluate the search with queries that are not indexed, we split the tiles of a dataset at random into two disjoint sets: the index tiles, from which the index is built, and the *held-out* tiles, which are used only as queries. Let *M* ⊆ {1, …, *M*} be the indices of the index tiles, *H* = {1, …, *M*}/*M* the indices of the held-out tiles, and *n* = | *M* | the number of index tiles. The split is made after tiling and after the covariance matrices are computed, so held-out tiles and indexed tiles are preprocessed alike. When Spindle indexes are stored and deployed for a dataset (rather than for evaluation), all tiles are indexed *M* = {1, …, *M* }.

#### B.2 Block distance

Spindle compares covariance matrices using the log-Euclidean distance [18, 19]. This distance expresses covariance comparison as Euclidean comparison of matrix logarithms, which allows us to construct vector representations and distance bounds for indexing.

##### Log-Euclidean distance

Let 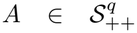 be a symmetric positive definite (SPD) matrix with eigendecomposition *A* = *V* diag(*λ*_1_, …, *λ*_*q*_)*V* ^⊺^, where *V* is orthogonal and *λ*_1_, …, *λ*_*q*_ *>* 0. The matrix logarithm of *A* is

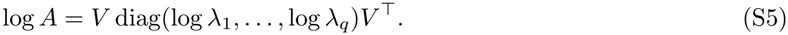

For two SPD matrices 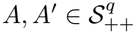, the log-Euclidean distance is

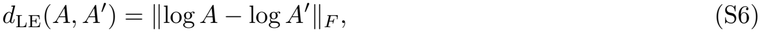

where ∥·∥_*F*_ denotes the Frobenius norm. The matrix logarithm is a bijection between SPD matrices and symmetric matrices. Thus, *d*_LE_ is a Euclidean distance in logarithmic coordinates and, in particular, is a metric satisfying the triangle inequality.

In numerical computations, we floor eigenvalues at 10^−6^ before taking their logarithms. This safeguard leaves Eq. (S5) unchanged for matrices whose eigenvalues are at least 10^−6^, including the regularized tile covariances defined above.

##### Block structures and block distance

Let *σ* be a permutation of the gene indices {1, …, *p*}, so that *σ*(1), …, *σ*(*p*) specifies the reordered genes in *G*. Given an SPD matrix *A*, Spindle considers a block structure *ℬ* = (*σ*; *B*_1_, …, *B*_*L*_), where *B*_1_, …, *B*_*L*_ form a partition of the gene indices into nonempty sets, each consisting of consecutive genes in the order specified by *σ*. Thus, *B*_*i*_ ∩ *B*_*j*_ = ∅ for *i* ≠ *j* and 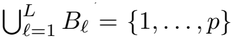. Let *p*_*ℓ*_ = |*B*_*ℓ*_|, so that 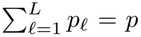, and let 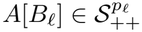 denote the corresponding principal submatrix, with its rows and columns ordered according to *σ*. In these reordered coordinates, the block-diagonal part of *A* under *ℬ* is

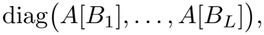

obtained by retaining entries within each block and setting all entries outside these blocks to zero. For notational convenience, we write *A*_*ℓ*_ for *A*[*B*_*ℓ*_].

###### Proposition S1

*Let A* = diag(*A*_1_, …, *A*_*L*_) *and A*′ = diag(*A*′1, …, *A*′*L*) *be block-diagonal SPD matrices with the same block sizes*, 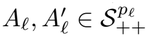. *Then* log *A* = diag(log *A*_1_, …, log *A*_*L*_), *and*

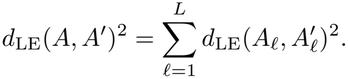

*Proof* Let *A*_*ℓ*_ = *V*_*ℓ*_Λ_*ℓ*_*Vℓ*⊺ be the eigendecomposition of block *ℓ*, with *V*_*ℓ*_ orthogonal and Λ_*ℓ*_ diagonal with positive entries. Then *V* = diag(*V*_1_, …, *V*_*L*_) is orthogonal, Λ = diag(Λ_1_, …, Λ_*L*_) is diagonal with positive entries, and *A* = *V* Λ*V* ^⊺^ is an eigendecomposition of *A*. Applying Eq. (S5) to this eigendecomposition gives log *A* = *V* log(Λ)*V* ^⊺^ = diag *V*_1_ log(Λ_1_)*V*1⊺, …, *V*_*L*_ log(Λ_*L*_)*VL*⊺ = diag(log *A*_1_, …, log *A*_*L*_), where log(Λ) denotes the diagonal matrix obtained by taking the logarithm of each diagonal entry.

The same holds for *A*′, so log *A* − log *A*′ = diag(log *A*_1_ − log *A*′1, …, log *A*_*L*_ − log *AL′* ) is block-diagonal. The squared Frobenius norm of a block-diagonal matrix is the sum of the squared Frobenius norms of its blocks, which gives the identity.

Let *ℬ* = (*σ*; *B*_1_, …, *B*_*L*_) be a block structure and *A*, 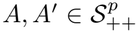 . We call log *A*[*B*_*ℓ*_], the matrix logarithm of the submatrix of *A* on the genes of block *B*_*ℓ*_, the *block logarithm* of *A* at block *ℓ*; Spindle-naive stores the block logarithms of every indexed tile (Section B.4). The distance between *A* and *A*′ at block *ℓ* is

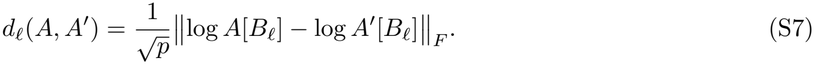

Consequently the block distance between *A* and *A*′ under *ℬ* is

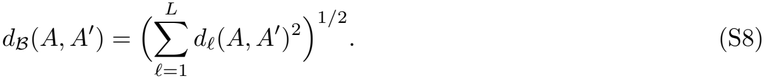

The factor 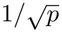 normalizes the squared distance by the number of genes, making the log-Euclidean distance comparable across datasets of different dimensions.

##### Partial block distance

Given a nonempty subset *S* ⊆ *G*, we define the partial block distance between *A* and *A*′ under *ℬ* as

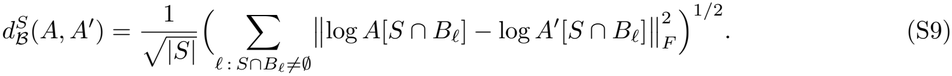

The sum includes only blocks that contain at least one gene in *S*. Spindle first extracts each submatrix and then takes its matrix logarithm. Taking the logarithm of the full block first and then extracting the submatrix generally gives a different result. Our definition requires covariance values only for genes in *S*, so it can be used when the query covariance is available only on those genes. When *S* = *G*, this definition reduces to *d*_*ℬ*_.

##### Block-log vectors

Given an SPD matrix 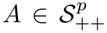 and a corresponding block structure *ℬ* = (*σ*; *B*_1_, …, *B*_*L*_), let 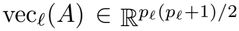 be the flattened vector of the upper triangle of log *A*[*B*_*ℓ*_] (Eq. (S5)) where diagonal entries are scaled by 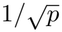 and off-diagonal entries are scaled by 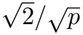. The block-log vector of *A* is

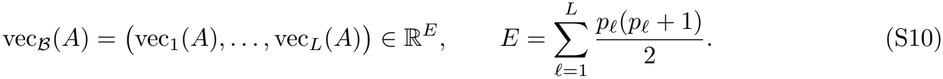

The scaling allows the lower triangle to be discarded without changing the distance:

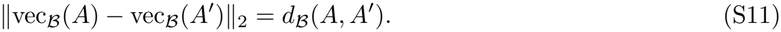

#### B.3 Covariance niches and block structures

Gene–gene covariance patterns vary across tissue regions, so a single block structure may not represent all tiles well. Spindle therefore groups index tiles with similar covariance organization into *covariance niches* and constructs a separate gene order and block partition for each niche. These structures are learned from index tiles only; every query is subsequently compared with tiles in every niche.

Spindle partitions the index tiles *M* into *J* disjoint covariance niches *N*_1_, …, *N*_*J*_ . Each niche *N*_*j*_ has a gene order *σ*_*j*_ and a block structure 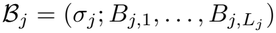. These determine how the query is compared with tiles in that niche. We describe their construction below; the index representations built on them are described in Sections B.4 and B.5.

The construction of the niches and block structures relies on two *modeling assumptions* :

1. **Latent niche structure**. There exists an embedding 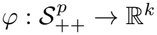, with *k* ≪ *p*(*p* + 1)*/*2, in which tile covariances form groups of similar co-variation patterns. These groups define the covariance niches.
2. **Consensus block structure within niches**. Each niche *N*_*j*_ admits a block structure *ℬ*_*j*_ under which its tile covariances are approximately block-diagonal.

By Proposition S1, the block distance equals the normalized log-Euclidean distance when both matrices are block-diagonal under the same block structure. This representation requires 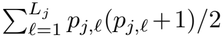 distinct block-log entries per tile in niche *j*, compared with *p*(*p* + 1)*/*2 for a full matrix logarithm. Given the indexed tile covariances {Σ_*m*_}_*m*∈*M*_, Spindle constructs the niches and their block structures in three steps: tile embedding, niche formation, and gene ordering and block selection within each niche.

##### B.3.1 Tile embedding

Spindle embeds tile covariances in a low-dimensional space to identify covariance niches (Assumption 1). For each indexed tile *m*, it first computes the *p* × *p* correlation matrix *R*^(*m*)^ from Σ_*m*_:

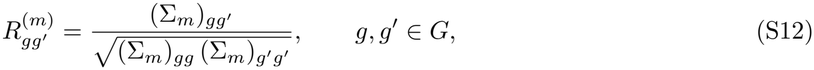

where (Σ_*m*_)_*gg′*_ denotes the covariance between genes *g* and *g*′. Spindle defines the gene dissimilarity as 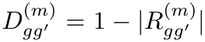 and constructs the dissimilarity matrix *D*^(*m*)^. This dissimilarity is zero on the diagonal and small for strongly correlated or anti-correlated gene pairs.

Spindle applies agglomerative hierarchical clustering with average linkage (UPGMA) to *D*^(*m*)^ [26]. The distance between two gene clusters *G*_1_ and *G*_2_ is

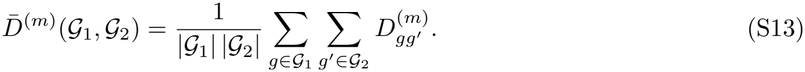

The resulting dendrogram defines the cophenetic distance 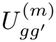 as the height at which genes *g* and *g*′ first merge into the same cluster [27]. The matrix *U* ^(*m*)^, with 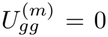, encodes the hierarchical relationships among genes in tile *m*.

Let *u*_*m*_ ∈ ℝ*p*(*p*−1)*/*2 contain the upper-triangular entries of *U* ^(*m*)^ in the reference gene order. Spindle applies PCA [38] to {*u*_*m*_}_*m*∈*M*_, with mean centering and no whitening, retaining *k* = min(30, *n* − 1) components. The resulting scores define the embedding *φ*(Σ_*m*_) ∈ ℝ*k* used to form niches. Queries are not embedded, as every niche is searched. The matrices *U* ^(*m*)^ are also used to derive each niche’s gene order.

##### B.3.2 Building covariance niches

Spindle constructs an undirected 10-nearest-neighbor graph of the tile embeddings (see Section B.3.1), using Euclidean distance Δ_*mm′*_ = ∥*φ*(Σ_*m*_) − *φ*(Σ_*m′*_ )∥_2_ and edge weights

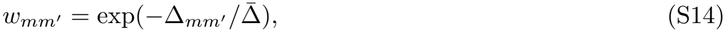

where 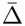 is the mean distance over all *n*^2^ ordered tile pairs.

The graph is partitioned using Leiden community detection [39] with modularity under the configuration null model. Spindle increases the resolution parameter *γ* from 0.2 to 1.6 in steps of 0.1, stopping when every community has at most *n*_max_ = 1,000 tiles. Any remaining oversized community *C* is split into ⌈|*C*| */n*_max_⌉ subcommunities using *k*-means clustering [48] on its tile embeddings. Splitting is repeated until every community satisfies the size limit.

##### B.3.3 Niche-specific block-diagonalization

###### Gene order within a niche

For each niche *N*_*j*_, Spindle averages the tile cophenetic matrices:

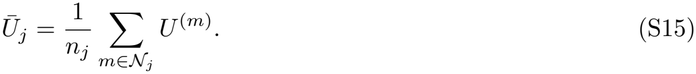

Spindle uses UPGMA clustering of 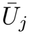 (Eq. (S13)) to create a consensus dendrogram *T*_*j*_ (Supplementary Fig. S8). Its left-to-right leaf order defines *σ*_*j*_. Spindle uses *σ*_*j*_ as the gene order shared by all tiles within niche *N*_*j*_.

For block selection, Spindle also computes the mean correlation matrix

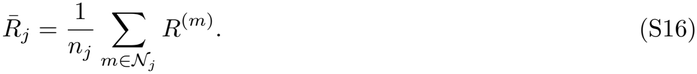

The problem of finding one gene permutation under which a set of matrices is approximately block-diagonal is known as approximate joint block diagonalization [49–51]. Its solvers optimize over orthogonal transformations of all matrices of the set, and their cost grows with the number of matrices. Spindle restricts the transformation to a permutation of the genes and obtains it from one consensus tree per niche, whose cost does not depend on the number of tiles beyond the averages of Eqs. (S15) and (S16), and which keeps every block a set of named genes.

###### Blocks within a niche

Spindle determines the blocks of niche *N*_*j*_ in two steps: it cuts the consensus dendrogram *T*_*j*_ into fine runs of consecutive genes, and then merges consecutive runs into blocks (Supplementary Fig. S8).

###### Step 1: Fine runs

Cutting the dendrogram *T*_*j*_ at height *t* induces a partition of the genes into runs. Let *F*(*t*) denote these runs in the order *σ*_*j*_, and let 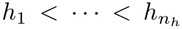 be the distinct merge heights of *T*_*j*_. We consider cuts at *t*_*r*_ = (*h*_*r*_ + *h*_*r*+1_)*/*2, for *r* = 1, …, *n*_*h*_ − 1. A cut is admissible if every run contains at most 50 genes.

If no cut is admissible, Spindle selects the candidate cut with the smallest maximum run size. Otherwise, let 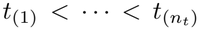 denote the admissible cut heights and define *ψ*_*i*_ = log |*F*(*t*_(*i*)_)|. If *n*_*t*_ *<* 3, Spindle selects *t*_(1)_. Otherwise, it smooths *ψ*_*i*_ using a nine-point moving average with zero padding and estimates the second derivative with respect to cut height by applying central finite differences twice. Let *i*^*^ minimize this estimated derivative. Spindle selects *t*_(max(*i*\*−2,1))_ as a heuristic knee of the run-count curve. In either case, the selected cut defines the fine runs *F*_1_, …, *F*_*R*_, ordered according to *σ*_*j*_.

###### Step 2: Merging runs into blocks

Given the fine runs *F*_1_, …, *F*_*R*_, Spindle selects boundaries

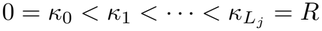

and forms blocks

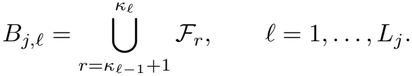

Thus, each block contains one or more consecutive fine runs, and no run is split.

For a candidate block *B*, define

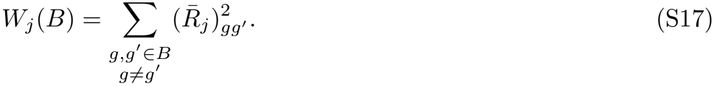

Spindle chooses the boundaries to maximize

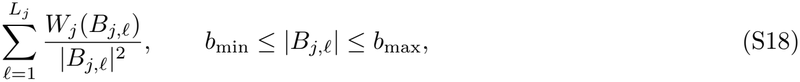

initially setting *b*_min_ = 15 and *b*_max_ = 100. The optimal boundaries are computed by dynamic programming over *F*_1_, …, *F*_*R*_, using two-dimensional prefix sums to evaluate *W*_*j*_. If no feasible partition exists, *b*_max_ is increased by 5 until one is found. The resulting blocks, together with *σ*_*j*_, define 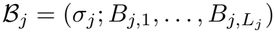.

#### B.4 Search problem and Spindle-naive

Given a query covariance *Q*, we search for similar tiles across all covariance niches. Let *j*(*m*) denote the unique niche index such that *m* ∈ *N*_*j*(*m*)_. Each tile is compared with *Q* using the gene order and block structure of its niche:

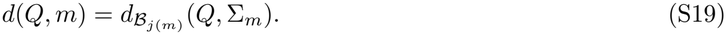

For a partial query *Q*_*S*_ over a nonempty gene subset *S* ⊆ *G*, we use

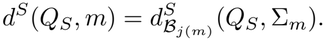

Using these distances, we define the budgeted covariance search problem as follows.

##### Problem S1

(**Budgeted covariance search**, Problem 1 restated). Given a query and a distance budget *η* ≥ 0, let *δ*(*m*) = *d*(*Q, m*) for a complete query or *δ*(*m*) = *d*^*S*^(*Q*_*S*_, *m*) for a partial query. Return a ranked list *ℛ* = (*m*_1_, …, *m*_*r*_) containing each tile in {*m* ∈ *M* : *δ*(*m*) ≤ *η*} exactly once, with

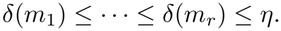

Ties are broken by tile number.

**Fig. S8:**
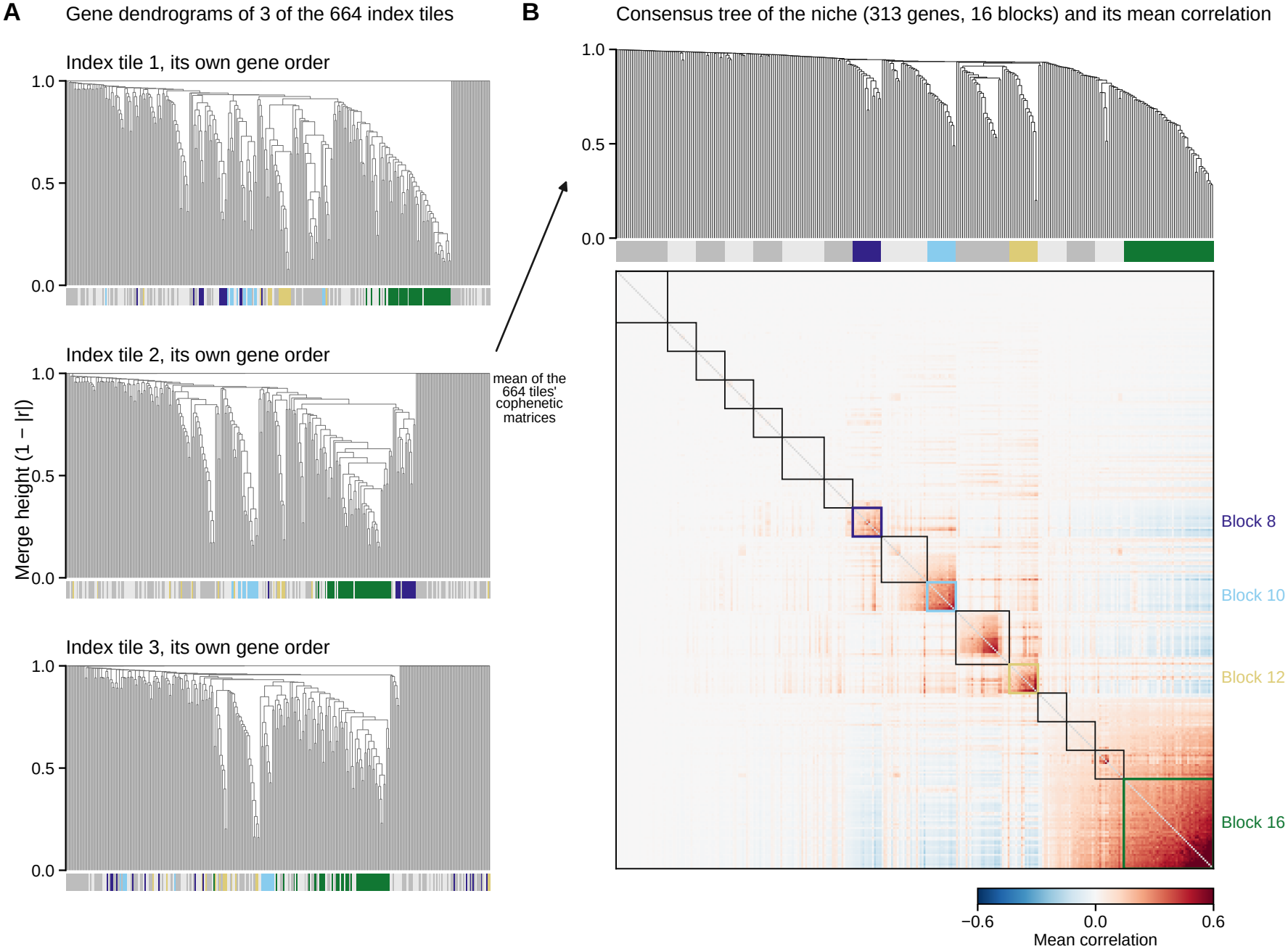
From tile dendrograms to the consensus tree and the blocks of the largest breast-cancer niche. **A** Gene dendrograms (average linkage on 1 − |*r*| ) of three randomly chosen index tiles of the niche, each in its own leaf order; the bar under the leaves colors each gene by its block in the niche. **B** The consensus tree, obtained by average linkage on the mean of the cophenetic matrices of the 664 index tiles of the niche; its leaf order is the niche’s gene order. Below, the 16 blocks (bar) and the mean correlation matrix of the niche in that order (diagonal not shown), with the blocks outlined. The four blocks that are subtrees of the consensus tree are colored and numbered; the other blocks are shown in alternating gray.

##### Exact exhaustive baseline

Spindle-naive solves Problem S1 by computing the distance to every indexed tile and returning those within the budget in increasing distance order. We first describe complete queries; the extension to partial queries is given in Section B.7.1.

For a niche *N*_*j*_ with *n*_*j*_ tiles, Spindle-naive stores the *n*_*j*_ × *E*_*j*_ matrix *Y*_*j*_ whose rows are the block-log vectors 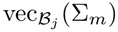 of the index tiles. The Spindle-naive index consists of the block structures *ℬ*_1_, …, *ℬ*_*J*_ and the matrices *Y*_1_, …, *Y*_*J*_ only. The logarithms of the index tiles are computed once, when the index is built. The index holds 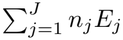 values, compared with *n p*(*p* + 1)*/*2 values for the logarithms of the whole covariance matrices.

Given a query covariance 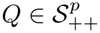, Spindle-naive computes its block-log vector under each niche’s block structure:

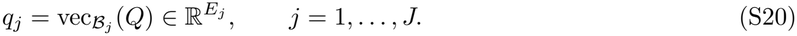

For each indexed tile *m* ∈ *N*_*j*_, the distance is computed using its stored block-log vector (Eqs. (S19) and (S11)):

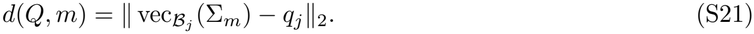

Scanning all indexed tiles requires Σ_*j*_ *n*_*j*_*E*_*j*_ subtractions and squares, with no matrix logarithms computed for indexed tiles. The tiles are returned with their distances in increasing order, with ties broken by tile index. For evaluations requesting a fixed number *c* of neighbors, we instead return the first *c* tiles of the exhaustive ranking.

#### B.5 Building the Spindle index

Instead of storing every tile’s block-log vector as Spindle-naive does, Spindle stores a representative set of block-log vectors for each block within each niche. A layered directed acyclic graph (DAG) connects these representatives across blocks: each layer corresponds to a block, and each tile follows a path through its assigned representatives. The DAG thus preserves the relationships between block representatives belonging to the same tile. Spindle uses the niche-specific block structures ℬ _*j*_ of Section B.3.

We describe the construction for a fixed niche *N*_*j*_, indexing its blocks by *b* and the DAG layers introduced below by *l*. For tile *m* ∈ *N*_*j*_, let

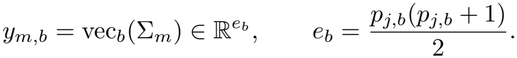

By Eq. (S11), ∥ *y*_*m,b*_ *y*_*m′,b*_ *∥* 2 = *d*_*b*_(Σ_*m*_, Σ_*m′*_ ).

Spindle repeats the construction for every niche.

##### DAG representation

Given a niche *N*_*j*_ whose tiles share the block structure 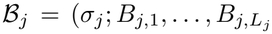), Spindle represents the niche by a layered DAG *G*_*j*_ = (*V, F*), where 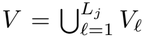 and *V*_*l*_ is the set of nodes in layer *l*. Each layer corresponds to one block, and each node *v* ∈ *V*_*l*_ stores a representative block-log vector *µ*_*v*_ and a radius *r*_*v*_.

Every tile *m* ∈ *N*_*j*_ is assigned to exactly one node in each layer. Let *M*_*v*_ be the set of tiles assigned to node *v*. The edge set is

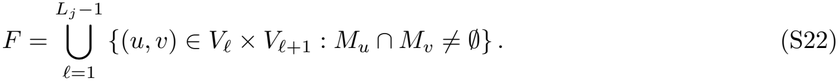

Edges connect only consecutive layers, so *G*_*j*_ is acyclic. Each tile follows a path through its assigned nodes, preserving the association between its blocks. For a path *P* = (*v*_1_, …, *v*_*s*_), the tiles following the entire path are

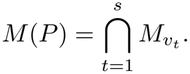

We next describe the layer order, the selection of representative vectors, and the tile assignments that determine the edges.

##### Layer order

The block assigned to each layer is determined by its spread across the tiles in *N*_*j*_. For block *B*_*j,b*_, define

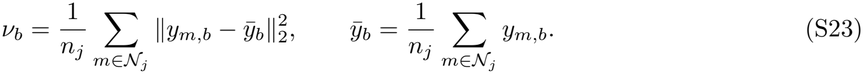

Let *π* order the blocks by decreasing spread, with ties broken by their order along *σ*_*j*_. Layer *V*_*l*_ represents block *B*_*j,π*(*l*)_, so

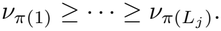

We write 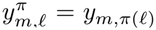 and 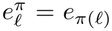_)_ for the block-log vector and its dimension in layer *l*. This ordering places the most variable blocks first to encourage early pruning as the search accumulates distance lower bounds across layers.

##### Representative nodes in a layer

For layer *V*_*l*_, we seek *K* representative tiles whose block-log vectors cover the layer with minimum radius. Let *ρ*_*l*_(*c*_1_, …, *c*_*K*_) denote the cover radius: the maximum distance from any tile’s block-log vector in layer *l* to its nearest center:

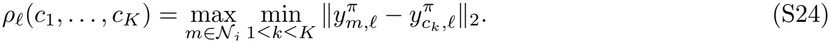

For each layer, we seek to solve the *K*-center problem defined below.

###### Problem S2

(*K*-center cover). Given the layer vectors 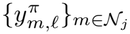 and 1 ≤ *K* ≤ *n*_*j*_, select *K* distinct center tiles *c*_1_, …, *c*_*K*_ ∈ *N*_*j*_ minimizing *ρ*_*l*_(*c*_1_, …, *c*_*K*_).

Spindle approximates Problem S2 using farthest-first traversal [40, 52]. Starting from one center, the algorithm repeatedly selects the tile farthest from its nearest existing center. For every prefix of *K* centers, the cover radius is at most twice [40] the optimal *K*-center radius. A single traversal therefore supplies candidate representatives for different layer sizes; the number retained in each layer is chosen later under a storage budget.

For each layer, the traversal starts with *c*_*l*,1_ = min *N*_*j*_ and selects at most min(*K*_max_, *n*_*j*_) centers, where *K*_max_ = 256. It stops earlier if the cover radius reaches zero (Algorithm S1). The output is a center sequence 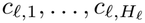 and its nonincreasing cover-radius curve

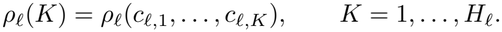

###### Algorithm S1

FarthestFirst({*y*_*m*_}_*m*∈*N*_, *K*_max_): center sequence and cover-radius curve of one layer

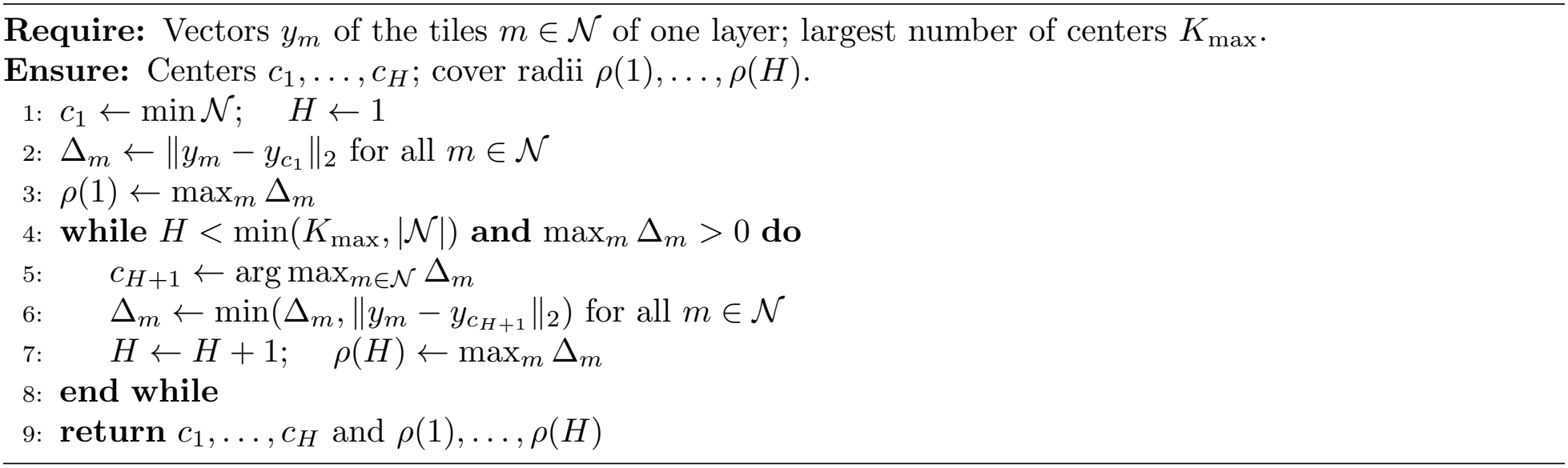

##### Node Allocation

Spindle retains the first *K*_*l*_ centers as the nodes *V*_*l*_ of layer *l*. It chooses *K*_*l*_ using a greedy allocation heuristic with a target storage budget of *βn*_*j*_*E*_*j*_ values, where *β* = 0.10, and 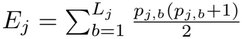 is the length of a tile’s block-log vector in niche *j*. Each node stores a center vector and a radius, requiring 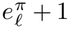 values. The budget excludes edges and tile-to-node assignments.

Initially, *K*_*l*_ = 1 for every layer. For a proposed increase to 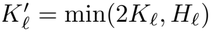, define

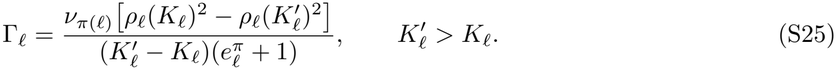

This score measures the reduction in squared cover radius, weighted by block spread, per additional stored value. At each step, the heuristic selects the layer *l* with the largest gain Γ_*l*_ and increases its number of retained centers from *K*_*l*_ to 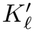 . It stops when node storage reaches the target budget or *K*_*l*_ = *H*_*l*_ for every layer.

The score balances representation error, block variability, and storage cost. Each *K*_*l*_ is a power of two or equals *H*_*l*_, and never exceeds *K*_max_.

##### Tile assignment and graph construction

After selecting *K*_*l*_ representatives for layer *V*_*l*_, Spindle assigns each tile to its nearest representative:

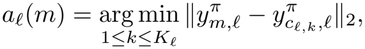

breaking ties by center index. Centers are indexed in their farthest-first selection order. Thus, *a*_*l*_(*m*) = *k* assigns tile *m* to the node (*l, k*) (*node index* ) represented by center tile *c*_*l,k*_.

For node *v* = (*l, k*), define

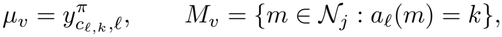

and

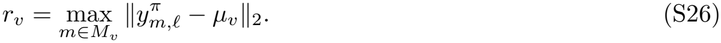

Edges are then determined by the shared-tile condition defined in Eq. (S22). Algorithm S2 summarizes the construction.

###### Algorithm S2

Construction of the Spindle index for niche N_*j*_

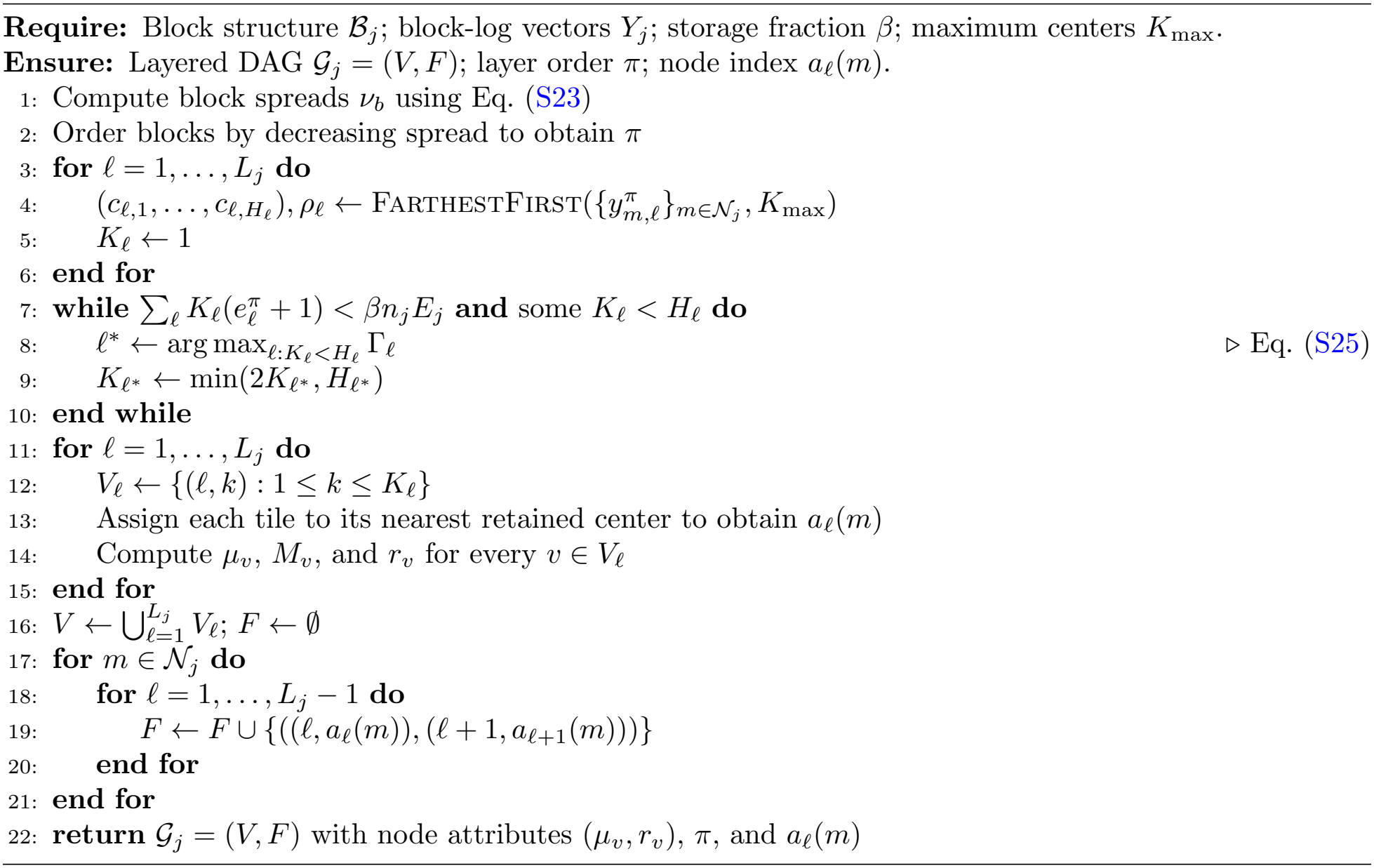

#### B.6 Searching Spindle

Spindle searches each niche’s DAG using node-level distance lower bounds to prune paths. A greedy probe sets a common distance budget across niches. The search returns a candidate set containing every tile within this budget and ranks the candidates using distances to their assigned representatives.

##### Query-to-node distances and lower bounds

For each niche *j*, Spindle computes the query vector *q*_*j*_ using Eq. (S20). Let 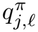 denote the query vector for layer *l*, containing the entries of *q*_*j*_ corresponding to block *π*(*l*). For a node *v* ∈ *V*_*l*_ with center *µ*_*v*_, define

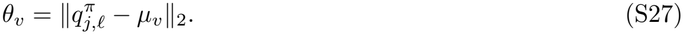

This distance is computed at most once per query and reused whenever the search revisits *v*.

For each node *v*, define

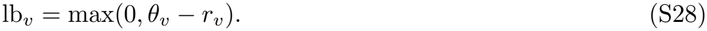

The following lemma shows how these values bound the distance of a query to a tile along its path.

###### Lemma S1.

*For a node v in layer l and any tile m* ∈ *M*_*v*_,

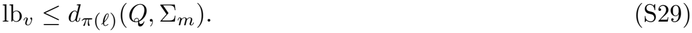

*Consequently, if* 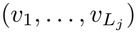 *is the path of tile m* ∈ *N*_*j*_, *then for every s* = 1, …, *L*_*j*_,

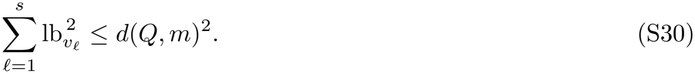

*Proof* For *m* ∈ *M*_*v*_, the triangle inequality gives

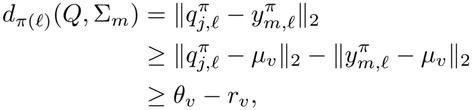

where the first equality follows from Eq. (S11) and the last inequality from Eq. (S26). Since *d*_*π*(*l*)_(*Q*, Σ_*m*_) 0, it follows that

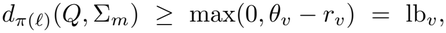

which is Eq. (S29). If 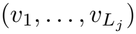 is the path of *m*, then 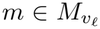 for every layer *l*, so squaring Eq. (S29) and summing over *l* ≤ *s* gives

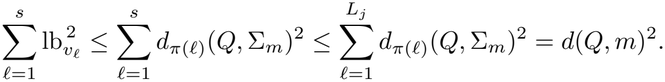

A tile *m* is reached by a path prefix *P*_*s*_ = (*v*_1_, …, *v*_*s*_) exactly when 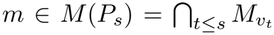, that is, when *P*_*s*_ is a prefix of its path.

##### Greedy probe and distance budget

Spindle selects a path greedily in each niche to set a common search budget. The probe starts at the node in the first layer with the smallest query-to-node distance *θ*_*v*_. At each subsequent layer, it extends the current path *P* to the child *v* minimizing *θ*_*v*_, subject to *M*(*P*) ∩ *M*_*v*_ ≠ ∅. This condition ensures that at least one tile is present in the complete path.

For the selected path 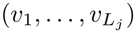 in niche *j*, define

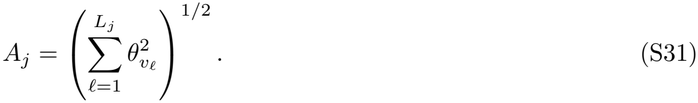

The search budget is

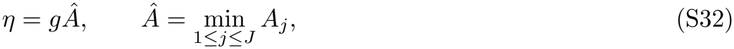

where *g >* 0 controls the search breadth and defaults to 0.25. The same budget is used in every niche. It scales with the smallest probe score rather than a fixed distance threshold.

##### Budgeted search and approximate ranking

Spindle searches every niche using the common budget *η* (Algorithm S3). For a path *P* = (*v*_1_, …, *v*_*s*_) starting in the first layer, define

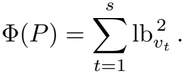

By Lemma S1, Φ(*P*) bounds the squared distance to every tile in *M*(*P*) from below. The search therefore prunes paths with Φ(*P*) *> η*^2^ and excludes extensions with no shared tile.

For each complete path reached, the search returns every tile *m* ∈ *M*(*P*) with approximate score

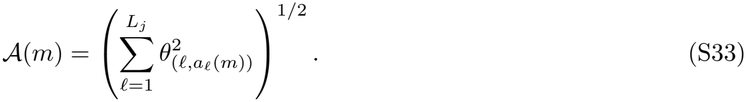

Results from all niches are ranked by *A*(*m*), with ties broken by niche and then tile index. The search imposes no additional limit on visited nodes or paths.

###### Theorem S2.

*For any g >* 0, *let η* = *gÂ be the budget defined in Eq*. (S32). *Algorithm S3 returns every indexed tile m satisfying d*(*Q, m*) ≤ *η*.

*Proof* Let *m* ∈ *N*_*j*_ satisfy *d*(*Q, m*) ≤ *η*, and let 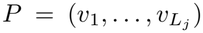 be its path. For every prefix *P*_*s*_ = (*v*_1_, …, *v*_*s*_), Lemma S1 gives

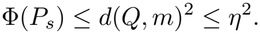

The first node *v*_1_ satisfies 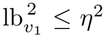. Since first-layer nodes are visited in increasing lower-bound order, *v*_1_ is visited before the stopping condition is met.

Suppose *P*_*s*_ is visited, where *s < L*_*j*_. Since 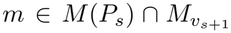, the next node *v*_*s*+1_ is a valid child. Moreover,

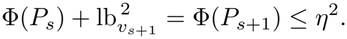

Children are considered in increasing lower-bound order, so the search visits *v*_*s*+1_ before stopping. Thus, *P*_*s*+1_ is visited. By induction, the search reaches *P* and returns *m* ∈ *M*(*P*). □

The theorem guarantees inclusion of every tile within the budget. The returned set may also contain tiles outside the budget, and its ordering by *A*(*m*) need not agree with the exact distance ordering.

###### Algorithm S3

SearchSpindle(*Q, g*): budgeted search of the Spindle index

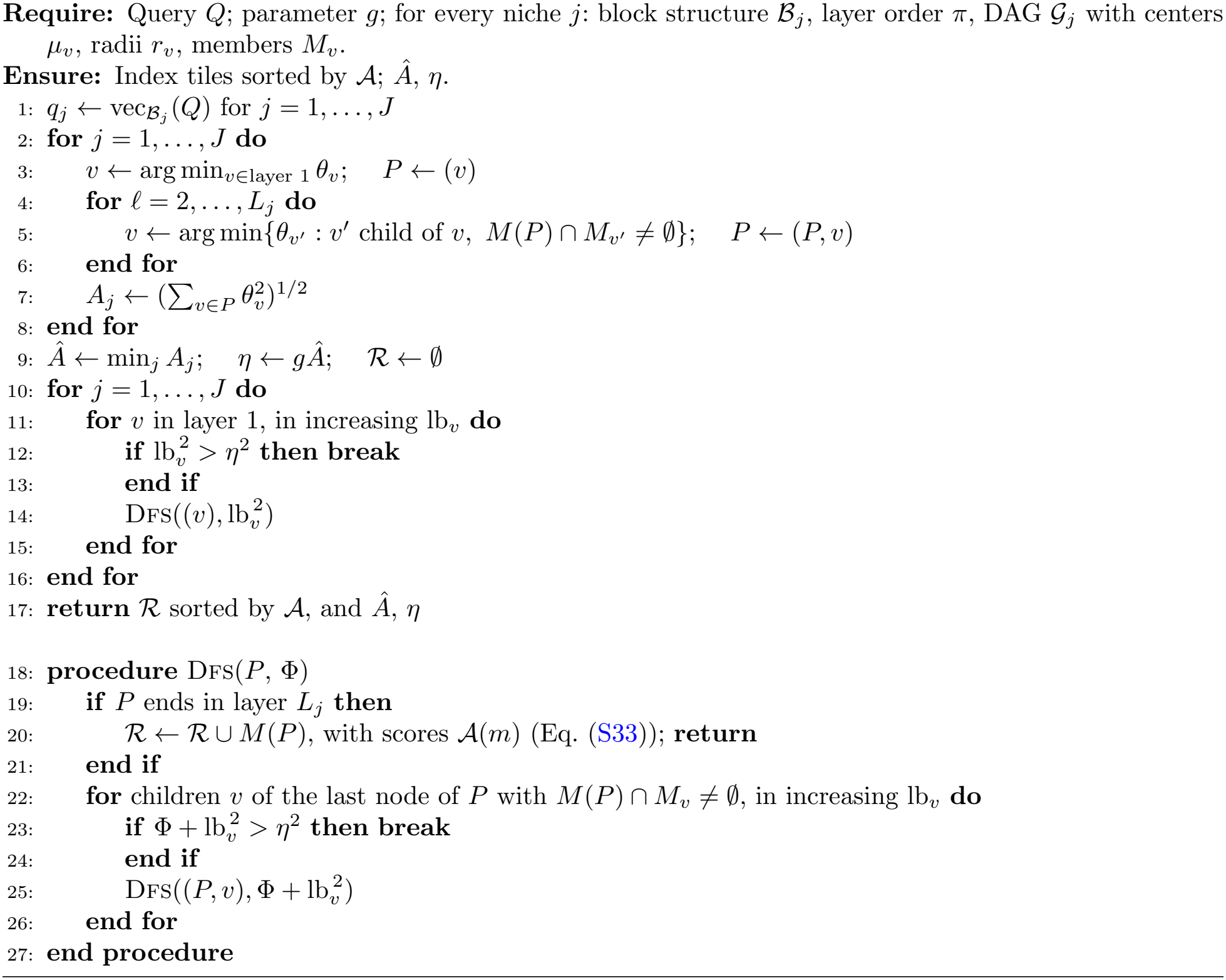

The compiled implementation performs the same search breadth first, layer by layer, discarding a tile when its accumulated lower bound exceeds *η*^2^; it returns the same tiles and scores as Algorithm S3.

#### B.7 Partial and gene-list queries

A partial query is a covariance matrix 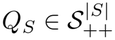 over a nonempty gene subset *S* ⊆ *G*. For example, *Q*_*S*_ may be the principal submatrix *Q*[*S*] of a complete query covariance. The search uses the partial block distance *d*^*S*^(*Q*_*S*_, *m*) defined in Section B.4.

To support partial queries, we extend Spindle-naive to evaluate the partial block distance exactly. We then construct Spindle-Partial using precomputed representations of dyadic intervals within each block. This interval decomposition introduces a finer block structure and a corresponding distance, defined below; the search guarantee applies to that distance. Both use the same niches and block structures as the indexes for complete queries.

##### B.7.1 Spindle-naive for partial queries

For exact partial-query search, Spindle-naive stores the covariance of each block in addition to its logarithm. For each niche *j* and block *B*_*j,b*_ intersecting *S*, Spindle-naive computes log *Q*_*S*_[*S* ∩ *B*_*j,b*_]. For each indexed tile *m* ∈ *N*_*j*_, it computes log Σ_*m*_[*S* ∩ *B*_*j,b*_] from the stored covariance, reusing the stored block logarithm when *S ∩ B*_*j,b*_ = *B*_*j,b*_. It then evaluates Eq. (S9) and returns tiles ranked by increasing distance, with ties broken by tile index.

##### B.7.2 Building Spindle-Partial

Following the convention of Section B.5, we describe the construction for a fixed niche *N*_*j*_. Spindle-Partial repeats the construction for every niche.

###### Dyadic intervals

Let *B*_*j,b*_ be a block of the niche with *p*_*b*_ genes. Let *g*_*b*_(*i*) denote the gene at position *i* in block *B*_*j,b*_, ordered according to *σ*_*j*_. For a set of positions *I*, define the corresponding gene set as *G*_*b*_(*I*) = {*g*_*b*_(*i*) : *i* ∈ *I*}. The covariance submatrix of tile *m* on these genes is Σ_*m*_[*G*_*b*_(*I*)].

###### Definition S1

(Dyadic interval). A dyadic interval in block *b* is a set of positions

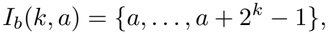

where *k, a* ≥ 0 are integers, *a* is a multiple of 2^*k*^, and *a* + 2^*k*^ ≤ *p*_*b*_. Let *I*_*b*_ denote all such intervals.

Positions are numbered from zero within each block. For each length 2^*k*^, there are ⌊*p*_*b*_*/*2^*k*^*⌋* disjoint dyadic intervals. Thus,

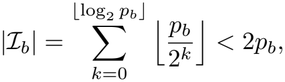

giving fewer than 2*p* intervals across all blocks of a niche. We order the intervals by block, then by increasing length and starting position.

###### Interval vectors and representative nodes

For each tile *m* ∈ *N*_*j*_ and interval *I* ∈ *I*_*b*_, let 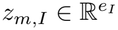 contain the upper-triangular entries of log Σ_*m*_[*G*_*b*_(*I*)], with off-diagonal entries multiplied by 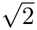, where

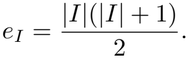

The covariance is restricted to *G*_*b*_(*I*) before taking its logarithm. This scaling preserves the log-Euclidean distance:

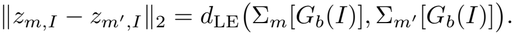

The vectors are stored without normalization by gene count; the search applies 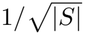 when combining interval distances. For each interval *I*, we apply the farthest-first traversal of Section B.5 to these vectors, obtaining *H*_*I*_ candidate centers and the cover-radius curve *ρ*_*I*_ (*K*). The allocation procedure below determines how many centers *K*_*I*_ to retain, giving the node set

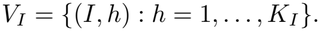

Each tile is assigned to its nearest retained center, with ties broken by center index. Let *a*_*I*_ (*m*) denote the assigned center index. Node centers *µ*_*v*_, membership sets *M*_*v*_, and radii *r*_*v*_ are defined as in the complete-query construction. The assignment table has rows ordered by tile index and columns ordered by interval. At query time, each selected interval contributes its node set as one search layer.

###### Node allocation

Following the node allocation procedure of section B.5, the numbers *K*_*I*_ are chosen by greedy doubling with a target node-storage budget of *βn*_*j*_*E*_*j*_ values, where *β* = 0.10 and *E*_*j*_ is the complete-query block-log vector dimension defined in Section B.5.

Each interval initially retains one center. Define its spread as

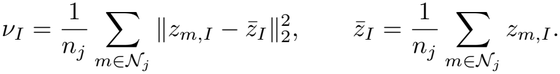

For a proposed increase from *K*_*I*_ to 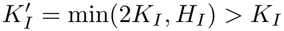, the gain is

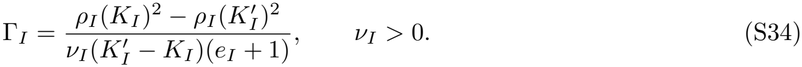

This measures the reduction in squared cover radius relative to interval spread, per additional stored value. At each step, the interval with the largest gain retains 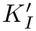 centers. The index stores ∑^*I*^ *K*_*I*_ (*e*_*I*_ + 1) values for centers and radii, and *n*_*j*_ ∑^*b*^ |*I*_*b*_| one-byte entries for tile-to-node assignments.

###### Traversing interval nodes

For a query on genes *S*, select an ordered sequence of dyadic intervals *I*_1_, …, *I*_*s*_ such that

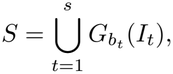

where *b*_*t*_ is the block containing *I*_*t*_. The interval-selection procedure is described below. These intervals define a layered DAG *G*_*j*_(*I*_1_, …, *I*_*s*_) = (*V, F*) with

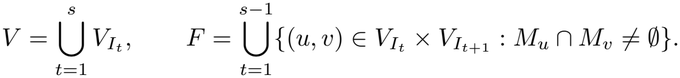

Each tile *m* follows the path

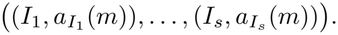

For a path *P* = (*v*_1_, …, *v*_*t*_), define

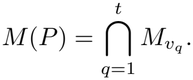

The nodes that can extend *P* while retaining at least one shared tile are

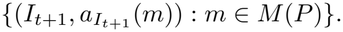

Thus, the node index determines the edges and eligible extensions for each query without storing a separate DAG for every interval sequence.

For selected intervals *I*_1_, …, *I*_*s*_, the node index determines connections between successive search layers. Nodes in consecutive layers are connected when they share an assigned tile. The search follows only connections that retain at least one tile common to the entire path.

##### B.7.3 Searching Spindle-Partial

Spindle-Partial uses Algorithm S3 for every niche with the modified DAG (Section B.7.2) *G*_*j*_(*I*_1_, …, *I*_*s*_) of the dyadic intervals.

###### Query decomposition

For each block *B*_*j,b*_, the positions of genes in *S ∩ B*_*j,b*_ are partitioned into maximal consecutive segments. Each segment [*a, e*) is decomposed into dyadic intervals from left to right using Algorithm S4. Starting at *c* = *a*, select *I*_*b*_(*k, c*) with the largest *k* satisfying 2^*k*^ |*c, c*+2^*k*^ ≤ *e*, and advance to *c*+2^*k*^ until the segment is covered. This produces a minimum-cardinality dyadic cover of the segment [53, 54] (Supplementary Fig. S9A).

**Fig. S9:**
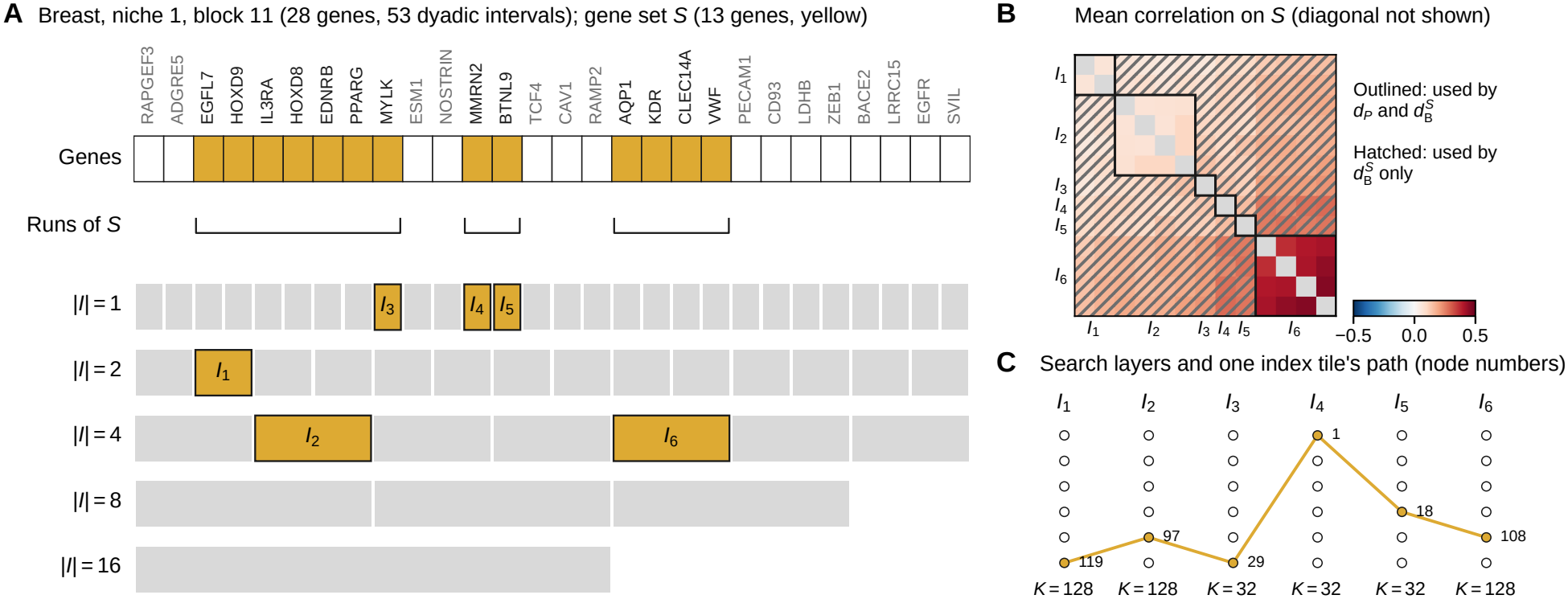
Dyadic decomposition of a partial query. **A** Block 11 of the largest breast-cancer niche (28 genes, in the niche’s gene order) and an illustrative gene set *S* of 13 genes (yellow). The maximal runs of *S* are covered, from left to right, by the fewest aligned dyadic intervals *I*_1_, …, *I*_6_ (yellow), shown among all 53 dyadic intervals of the block by length |*I*| (gray). **B** Mean correlation matrix of the niche on *S* (diagonal not shown): dP uses the entries within each piece (outlined), whereas 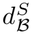 also uses the entries between pieces of the same block (hatched). **C** The pieces as the layers of the Spindle-Partial search, with the number of nodes *K*_*I*_ of each interval, and the path of one index tile through its nodes.

###### Algorithm S4

Dyadic decomposition of a segment [*a, e*) in block *b*

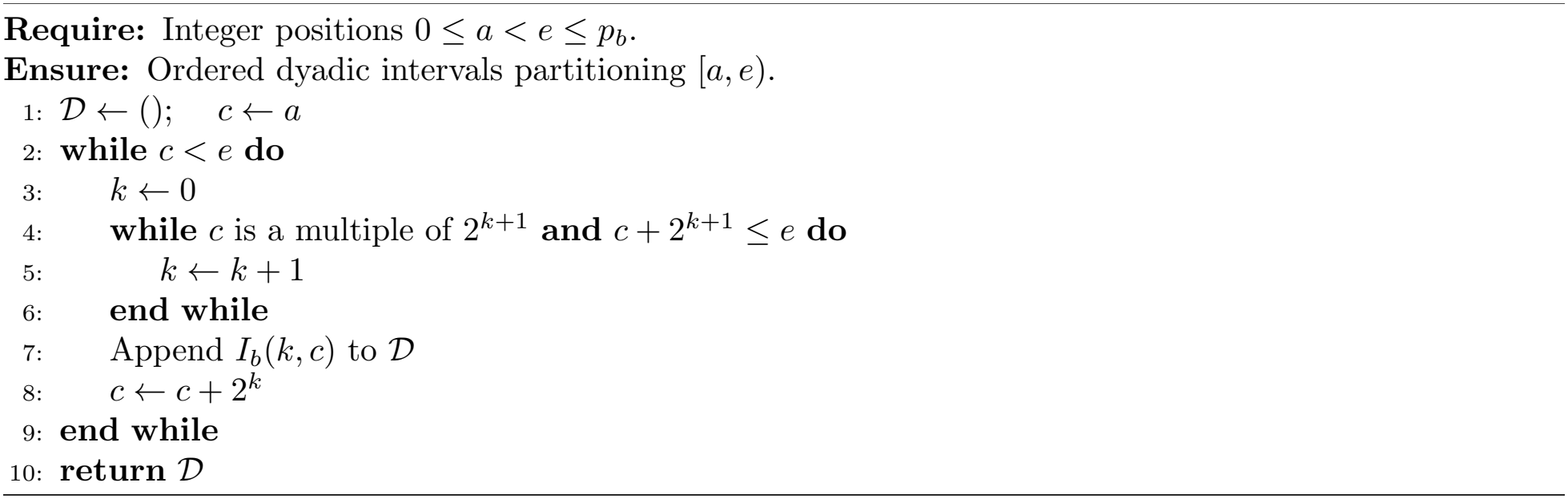

Within niche *j*, let *I*_1_, …, *I*_*s*_ be the selected intervals in block and position order, with *I*_*t*_ belonging to block *b*_*t*_. Their gene sets 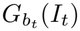 partition *S*. This decomposition depends on the niche’s gene order and block structure.

For each *I*_*t*_, we vectorize log 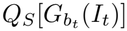 using the same scaling as 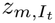 to obtain the query vector *w*_*t*_. The search traverses the corresponding node sets 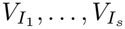 as successive DAG layers.

###### Distance over selected intervals

For a tile *m* ∈ *N*_*j*_, define

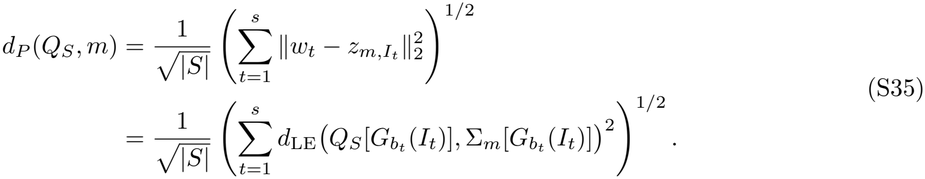

By Proposition S1, this is the normalized log-Euclidean distance under the finer block structure defined by the selected intervals. Compared with *d*^*S*^(*Q*_*S*_, *m*), it additionally omits covariances between different intervals within the same block. If each nonempty intersection *S* ⋂ *B*_*j,b*_ is covered by a single dyadic interval, then *d*_*P*_ (*Q*_*S*_, *m*) = *d*^*S*^(*Q*_*S*_, *m*).

###### Budgeted search

For each niche *j*, Algorithm S3 traverses the node sets 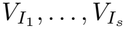 in the selected interval order. For a node 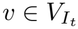, define

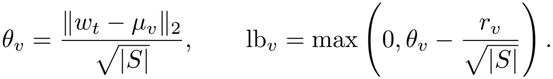

These quantities use the same normalization as *d*_*P*_ . Tile assignments in the node index determine eligible path extensions, as described in the construction.

The greedy probe, budget *η* = *gÂ*, path bound Φ(*P*), and ranking score *A*(*m*) are computed using these node distances, with *g* = 1 by default. For every tile *m* and every prefix *P*_*t*_ of its path, the argument of Lemma S1 gives

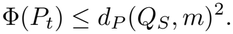

Thus, the same pruning guarantee applies.

###### Theorem S3.

*For any g >* 0, *Algorithm S3, applied to the selected interval layers with the node distances above, returns every indexed tile m satisfying d*_*P*_ (*Q*_*S*_, *m*) ≤ *η, where η* = *gÂ is the budget obtained from the partial-query greedy probes*.

*Proof* For any tile *m* satisfying *d*_*P*_ (*Q*_*S*_, *m*) ≤ *η*, every prefix *P*_*t*_ of its path contains *m* in *M*(*P*_*t*_) and satisfies Φ(*P*_*t*_) ≤ *η*^2^. Hence the induction in the proof of Theorem S2 applies. □

This guarantee concerns *d*_*P*_, rather than *d*^*S*^; the two coincide when each nonempty block intersection is represented by a single selected interval.

###### Gene-list queries

When only a gene set *S* is provided, Spindle constructs *Q*_*S*_ using a heuristic co-expression template:

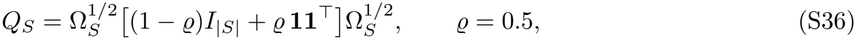

where *I*_|*S*|_ is the |*S*| × |*S*| identity matrix, **1** is the all-ones vector of length |*S*|, and Ω_*S*_ is the |*S*| × |*S*| diagonal matrix whose entry for gene *g* is the 90th percentile of the variance (Σ_*m*_)_*gg*_ of gene *g* over the index tiles *m* ∈ *M*.

#### B.8 Implementation

Spindle is implemented in Python with NumPy [55] and SciPy [56], and builds the covariance niches with igraph [57] and the Leiden algorithm [39]; the searches of the Spindle-naive, Spindle and Spindle-Partial indexes run in compiled C kernels that return the same results as the Python reference implementation. All experiments ran on one compute node under Ubuntu 24.04 through the Slurm scheduler, without GPUs. Every query ran on one core, except in the multi-core timing (section A.5), where the search of Spindle processed the niches of a query on separate cores and returned the same tiles and scores as on one core.

